# Coronavirus protein interaction mapping in bat and human cells identifies molecular and genetic switches for immune evasion and replication

**DOI:** 10.1101/2025.07.26.666918

**Authors:** Jyoti Batra, Magdalena Rutkowska, Yuan Zhou, Chengjin Ye, Rithika Adavikolanu, Janet M. Young, Durga Anand, Sooraj Verma, Martin Gordon, Shivali Malpotra, Anastasija Cupic, Thomas Kehrer, Jack M Moen, Declan M. Winters, Ajda Rojc, Ignacio Mena, Sadaf Aslam, Carles Martinez-Romero, Isabela Conde Viñas, Zain Khalil, Keith Farrugia, Atoshi Banerjee, Dafna Tussia-Cohen, Melanie Dos Santos, Sourobh Maji, Monita Muralidharan, Helene Foussard, Irene P. Chen, Rotem Fuchs, CJ San Felipe, Lorena Zuliani-Alvarez, Promisree Choudhury, Kirsten Obernier, Ségolène Gracias, Rahul Suryawanshi, Carlos Ibáñez, Javier Juste, Lars Pache, Taha Y. Taha, Nolwenn Jouvenet, Kliment A. Verba, James S Fraser, Caroline Demeret, Robert M. Stroud, Harm van Bakel, Melanie Ott, Tzachi Hagai, Ben Polacco, Danielle L Swaney, Ignacia Echeverria, Mehdi Bouhaddou, Manon Eckhardt, Harmit S. Malik, Luis Martinez-Sobrido, Lisa Miorin, Adolfo García-Sastre, Nevan J Krogan

## Abstract

Coronaviruses, including SARS-CoV-2, can cause severe disease in humans, whereas reservoir hosts like *Rhinolophus* bats remain asymptomatic. To investigate how host-specific protein-protein interactions (PPIs) influence infection, we generated comparative PPI maps for SARS-CoV-2 and its bat-origin relative RaTG13 using affinity purification-mass spectrometry (AP-MS) in human and *Rhinolophus ferrumequinum* (RFe) bat cells. This approach identified both conserved and virus- and host-specific interactions that regulate infection dynamics. Notably, SARS-CoV-2 required a non-synonymous mutation in nucleocapsid to replicate in bat cells expressing human ACE2 and TMPRSS2. Analysis of the viral protein Orf9b revealed differential interactions with mitochondrial proteins Tom70 and MTARC2. A single residue difference in Orf9b between SARS-CoV-2 and RaTG13 functions as a molecular switch, weakening Tom70 binding and immune evasion in human cells while enhancing interaction with the bat-specific restriction factor MTARC2. These findings demonstrate how a single-residue substitution can reshape virus-host interactions and contribute to immune evasion and host adaptation.

## Introduction

Severe acute respiratory syndrome coronavirus 2 (SARS-CoV-2) is the causative agent of the COVID-19 pandemic, resulting in over 770 million cases and over 7 million deaths globally (source: WHO^1^). SARS-CoV- 2-related viruses (SARSr-CoVs) have been identified in *Rhinolophus* bats^2^ and pangolins^3^, although the definitive animal reservoir of SARS-CoV-2 remains uncertain. *Rhinolophus* bats (also called horseshoe bats) are considered key natural hosts of SARS-related coronaviruses (SARSr-CoVs), which are classified within the sarbecovirus subgenus, including SARS-CoV, which caused the 2003 outbreak^4^. Their prevalence as SARSr-CoV reservoirs is likely due to a unique balance between antiviral defense mechanisms and immune tolerance in bats. For example, some bat species have constitutive expression of interferon and interferon stimulated genes (ISGs), elevated heat shock proteins (HSPs), dampened STING activity, suppressed NLRP3 inflammasome pathways, and reduced IL-1β expression; all these factors likely contribute to making bats potentially asymptomatic and tolerant viral hosts^5^.

*Rhinolophus* bats are distributed across Asia, Southern Europe, and North Africa but are absent in the Americas. In Europe, SARSr-CoVs have been detected in *R. ferrumequinum*, *R. hipposideros*, *R. euryale*, and *R. mehelyi*^6–8^, whereas *R. sinicus*, *R. ferrumequinum*, and *R. affinis* are primary reservoirs in Asia, with seropositivity rates ranging up to 60%^9^. Among the closest known relatives of SARS-CoV-2 are RaTG13, sampled from *R. affinis*^10^, and BANAL-52, isolated from *R. malayanus* in Laos^2^. RaTG13 shares 96.2% identity with SARS-CoV-2 throughout the length of the genome, while BANAL-52 exhibits a slightly higher overall genome identity (96.8%) with SARS-CoV-2, especially in the receptor-binding domain (RBD) of the spike gene^2^. This might explain BANAL-52’s higher affinity to hACE2 and increased ability to mediate virus entry into human cell lines in pseudovirus entry assays compared to RaTG13^2^. Despite the high prevalence of SARS-CoV-2-related viruses in bats, the molecular determinants of cross-species transmission and the mechanisms that enable bats to remain asymptomatic carriers in the absence of apparent disease symptoms remain poorly understood. Given the importance of host-virus interactions in facilitating or restricting viral transmission, studying these mechanisms in bat reservoir species is essential for a deeper understanding of coronavirus biology and zoonotic spillover events. And yet, very few of the several available bat cell lines support productive SARS-CoV-2 infection^11–14^, posing a significant barrier to studying mechanisms facilitating SARS-CoV-2 spillover events. Here we report the establishment of a fibroblast cell line derived from lung samples of *R. ferrumequinum* bats (RFe cell line). To directly investigate how these distinct host interactions impact viral infection, we established an infection model by engineering RFe cells to express hACE2 and hTMPRSS2 (RFe-AT), and selecting for a SARS-CoV-2 variant capable of efficient replication in these bat cells. This process identified a single amino acid substitution of proline to threonine in the N-terminal domain of the nucleocapsid protein (N_P80T) that is sufficient to enable productive viral replication in bat cells expressing human ACE2 and TMPRSS2. Our bat cellular infection model will be a valuable tool for investigating the evolutionary relationship between coronaviruses and their hosts, as well as to better understand the unique bat host responses to virus infection.

Since successful viral replication depends on complex protein-protein interactions (PPIs) between viral and host proteins, we further reasoned that comparative interactome analyses in bat and human cells could reveal host-specific factors that govern viral replication and immune evasion. Indeed, previous efforts to map PPIs between SARS-CoV-2 proteins and human proteins have delivered deep mechanistic insights into host responses to infection^15–17^. Thus, identifying both conserved and divergent interactions across species could uncover cellular pathways critical for viral infection, adaptation and zoonotic transmission, thereby improving our understanding of coronavirus biology and informing strategies to predict and mitigate future spillover events. Using affinity tag-purification coupled with mass spectrometry (AP-MS), we conducted comparative protein interaction mapping by expressing viral proteins from both RaTG13 and SARS-CoV-2 in RFe and human HEK293T cells. We mapped species-specific interaction networks, identifying several conserved interactions across host species that suggest critical roles for these host proteins in viral infection. Our previous findings showed that SARS-CoV-2 Orf9b, an accessory protein translated from an alternative open reading frame within the N gene, is implicated in innate immune evasion by targeting mitochondrial protein Tom70. In contrast, RaTG13 Orf9b exhibited reduced interaction with Tom70, resulting in diminished suppression of the innate immune response. We investigated the functional role of the Orf9b-Tom70 interaction by generating a recombinant SARS-CoV-2 carrying a mutation in Orf9b that disrupts binding to Tom70. This mutant virus was attenuated in both bat and human cells, leading to a heightened immune response. Conversely, RaTG13 Orf9b demonstrated stronger binding to another mitochondrial protein, MTARC2, specifically in bat cells. We showed that MTARC2 is a bat-specific restriction factor that suppresses viral replication, highlighting species-specific differences in antiviral defenses. Our findings thus reveal distinct mechanisms of immune modulation and viral persistence between human and bat hosts. Additionally, we identified several unique interactions in bat cells that suggest other distinct mechanisms underlying immune modulation and viral persistence in human versus bat cells. Overall, our findings enhance our understanding of how SARS-CoV-2 and related viruses adapt to different host environments and evade immune responses; such information is crucial for developing effective strategies to combat these pathogens.

## Results

### Comparative interactome profiling of SARS-CoV-2 and RaTG13 in bat and human cells reveals conserved and virus-specific interactions

Bats serve as natural reservoir hosts for a wide range of emerging viruses, including SARS-related coronaviruses (SARSr-CoVs), yet the molecular mechanisms that govern their unique ability to tolerate viral infections remain poorly understood. To investigate these mechanisms, we sought to develop an *in vitro* model to study host-virus interactions directly in bat cells and to characterize the cellular response to SARS-CoV-2 infection in a bat-host context. To date, most SARS-CoV-2 related viruses have been identified in *Rhinolophus* bats. Therefore, we focused on developing a SARS-CoV-2 infection model using cell lines derived from lung tissue of a *R. ferrumequinum* bat captured near Jerez de la Frontera, Cádiz, in southern Spain in 2020. Primary fibroblasts were first established from adult male bat lung tissue, as described in detail in the methods section. Briefly, tissue was enzymatically and mechanically dissociated, and cells were cultured to allow outgrowth. Next, cells were expanded and immortalized by nucleofection of a SV40 large T antigen (SV40-LT) expressing plasmid. These immortalized bat lung fibroblast cells are referred to as RFe cells throughout this study. To confirm the species origin of the cell line, we compared the cytochrome c and cytochrome b gene sequences obtained from bulk RNA sequencing of total RNA isolated from RFe cells to published NCBI Reference Sequences^18^, confirming >99.5% identity (**Table S1 and S2**). By abundance proteomics, we identified common lung fibroblast markers, including S100 calcium-binding protein A4 (S100A4), vimentin (VIM), and several proteins from the collagen family, which were more prominently detected in both RFe and the well-characterized human lung fibroblast cell line MRC5 cells compared to HEK293T, as expected (**Figure S1A**). Additionally, similar to MRC5 cells, the immortalized RFe cells displayed typical fibroblast morphology, including large, flat, spindle-shaped cells with oval nuclei (**Figure S1B**).

Leveraging these cell lines and extending our previous work on the SARS-CoV-2 interactome in HEK293T cells^17^, we systematically mapped host factors that physically associate with individual proteins from SARS-CoV-2 and RaTG13 in both RFe (bat) and HEK293T (human) cells. SARS-CoV-2 and RaTG13 share high sequence similarity, with several viral proteins being nearly identical, including eight (Nsp7-11, Nsp16, E and Orf6) being 100% identical at the amino acid level (**Figure S2A**). For interactome mapping, we codon-optimized sequences encoding structural proteins, processed nonstructural proteins (Nsps), and predicted open reading frame (Orf) proteins and cloned them into mammalian expression vectors, pLVX-EF1a and pLVX-TetOne, incorporating a 2X-Strep tag for efficient purification. The pLVX-EF1a viral constructs were expressed in HEK293T cells by transient transfection. In contrast, due to low transfection efficiency in bat cells, we utilized the doxycycline-inducible pLVX-TetOne system to generate stable cell lines for protein expression. Each viral protein was affinity purified, and interacting host factors were identified using mass spectrometry (MS). High-confidence interactions were determined using SAINTexpress (Significance Analysis of Interactome)^19^ and MiST (Mass Spectrometry Interaction Statistics)^20^ scoring algorithms (**Figure 1A**).

**Figure 1.**
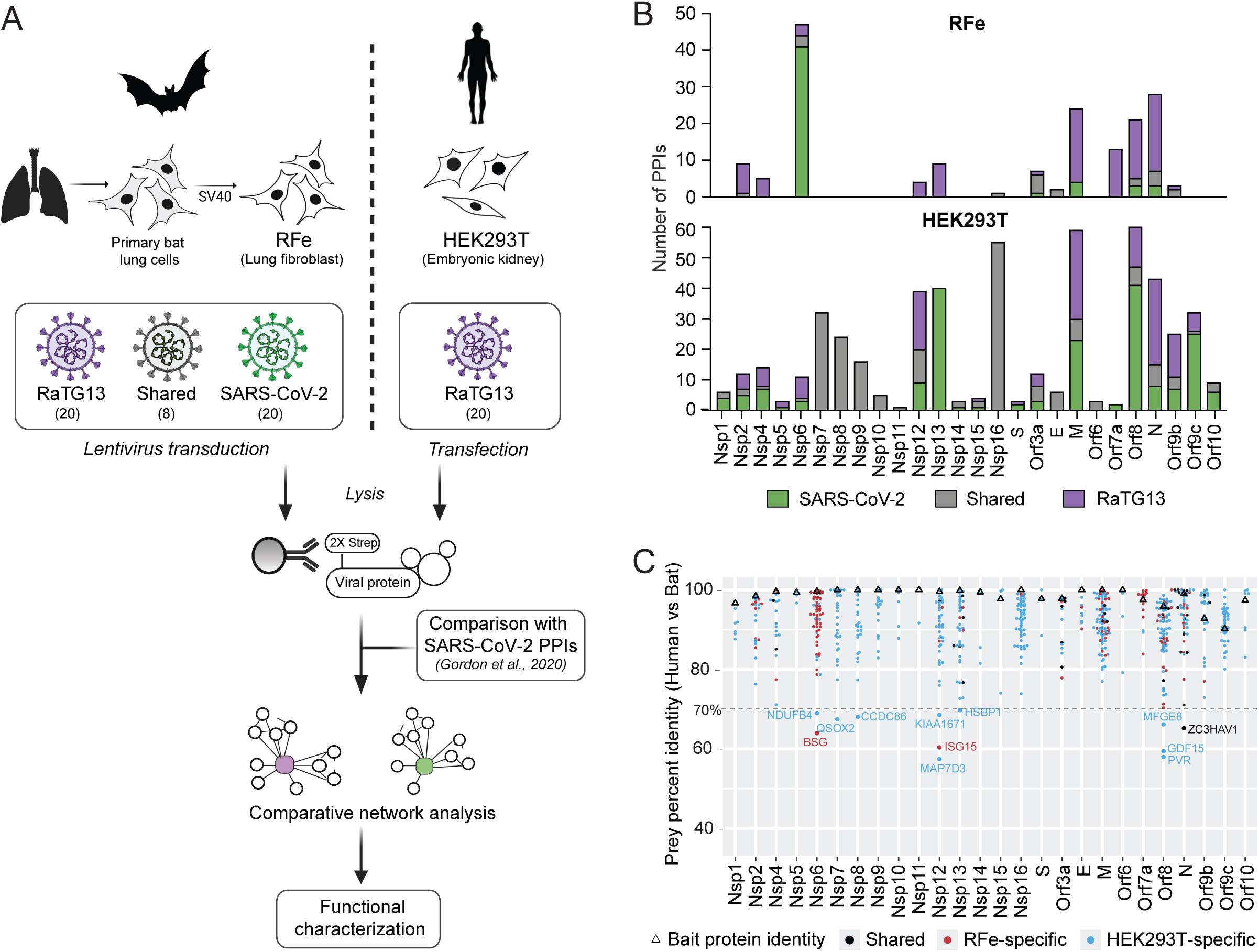
An affinity purification coupled to mass spectrometry (AP-MS) approach for comparative network analysis of SARSr-CoVs. (A) Schematic overview of the AP-MS workflow for mapping protein-protein interactions (PPIs) of SARS-CoV-2 and RaTG13 in human (HEK293T) and bat (RFe) cells. Primary lung cells from *Rhinolophus ferrumequinum* were immortalized using SV40 large T antigen to generate RFe lung fibroblast cells, which were used for proteomic studies. Viral proteins from SARS-CoV-2 and RaTG13 (n=20 per virus), along with 8 proteins that are identical between the two viruses, were tagged with 2X-Strep and expressed in RFe cells upon lentiviral transduction under a doxycycline-inducible promoter. Expression was induced for 24 hours prior to cell lysis. In parallel, HEK293T cells were transiently transfected with the 2X-Strep-tagged viral constructs cloned into the pLVX-EF1α vector. Strep-tagged viral proteins were affinity-purified, and co-purifying host proteins were identified by mass spectrometry. Comparative network analyses were performed, including integration with previously published SARS-CoV-2 PPIs (Gordon et al., 2020), to identify conserved and cell-type/virus-specific interactions. (B) Bar plot showing the number of high-confidence PPIs identified for each viral protein of SARS-CoV-2 and RaTG13 in RFe (top) and HEK293T (bottom) cells. Interactions are color-coded as SARS-CoV-2-specific (green), RaTG13-specific (purple), or shared (gray). (C) Dot plot showing the percent protein sequence identity of all high-confidence prey proteins between humans and *R. ferrumequinum*, grouped by their corresponding viral bait protein. Each dot represents a prey protein detected in AP-MS experiments with either RaTG13 or SARS-CoV-2 viral baits. Prey identities were determined by aligning human and *R. ferrumequinum* orthologs, and percent identity was calculated. Shared interactors (detected in both cell types) are shown in black; RFe-specific interactors in red; and HEK293T- specific interactors in blue. Viral bait protein identities (between SARS-CoV-2 and RaTG13) are indicated by open triangles.

Using this approach, we identified 72 high-confidence PPIs of SARS-CoV-2 in RFe cells, as well as 118 and 189 interactors of RaTG13 in RFe and HEK293T cells, respectively (**Figure 1B, Table S3)**. Previously, we described 332 high confidence PPIs of SARS-CoV-2 in HEK293T cells^17^. Among the 28 bait proteins, Orf3b and Orf7b showed an unusually high number of background PPIs and were therefore excluded from protein interaction analysis. Notably, Nsp6 and Orf7a showed a higher number of PPIs in RFe cells compared to HEK293T cells. We observed fewer SARS-CoV-2 PPIs compared to RaTG13 in bat cells, likely because several SARS-CoV-2 viral baits were expressed at significantly lower levels, as evidenced by log₂ intensity values and western blot (**Figure S2B-E**), likely reducing host protein enrichment and the number of detectable PPIs.

Despite the high similarity across the two viral isolates, we identified distinct interactomes for SARS-CoV-2 and RaTG13, with only 14% and 10% shared high-confidence interactions in HEK293T and RFe cells, respectively **(Figure S2F**). For RaTG13, ∼9% of interactions were conserved across both bat and human cells. To further investigate species-specific differences, we analyzed the sequence identity of host proteins interacting with SARS-CoV-2 and/or RaTG13 in bat or human cells. While most interacting host proteins displayed high sequence identity between bat and human orthologs, a subset of species-specific interactions involved prey proteins with lower sequence identity, including interactors of Orf8 and Nsp12 (**Figure 1C, Table S4**). These divergent orthologs may underlie functional differences in how viral proteins engage host pathways in bats versus humans, potentially shaping species-specific outcomes of infection and immune modulation. Notably, ZC3HAV1 (also known as ZAP) stood out as a shared interactor with RaTG13 N in both species. ZAP is a well-characterized antiviral effector that binds viral RNAs and promotes their degradation^21^. ZAP evolves rapidly in primates^22^ and has low sequence identity between the human and bat orthologs. Thus, even though the interaction between ZAP and RaTG13 N is conserved, sequence divergence in ZAP may influence species-specific differences in antiviral activity.

To quantitatively compare PPIs between SARS-CoV-2 and its close bat coronavirus relative RaTG13, we applied our previously developed differential interaction scoring (DIS) method^23^. This approach allowed us to identify conserved and virus-specific interactions for SARS-CoV-2 and RaTG13 across HEK293T and RFe cells (**Figure 2, S3**). The comparative interactome maps revealed a conserved set of host interactions, along with virus- and host-specific differences indicative of evolutionary adaptation and differential targeting of host pathways. In both host systems, viral proteins targeted key cellular pathways including RNA processing (Nop56p-associated pre-rRNA complex), translation (regulation of gene expression), trafficking (nuclear/mitochondrial transport), and innate immune regulation (Tom70, stress granule regulation). For example, the nucleocapsid (N) proteins of both viruses interact with the Nop56p-associated pre-rRNA complex (**Figure S3**), suggesting a conserved role in modulating host gene expression in both bat and human cells. Similarly, the accessory protein Orf8 of both viruses interacted with factors involved in ER organization and protein quality control in both species, consistent with prior reports showing Orf8 accumulation in the ER and its role in ER remodeling and homeostasis^24^. We also noticed several virus-specific differences. For example, in human cells, SARS-CoV-2 Nsp4 showed interaction with the mitochondrial translocase of the inner membrane (TIM) complex, while RaTG13-specific interactions were observed for Nsp12 with components of the MTOC (microtubule organizing center) (**Figure 2B**). We also observed a host species-specific interaction between RaTG13 Nsp2 and flotillin proteins (FLOT1 and FLOT2) in bat cells, indicating potential divergence in the targeting of membrane-associated trafficking pathways across species (**Figure S3A**).

**Figure 2.**
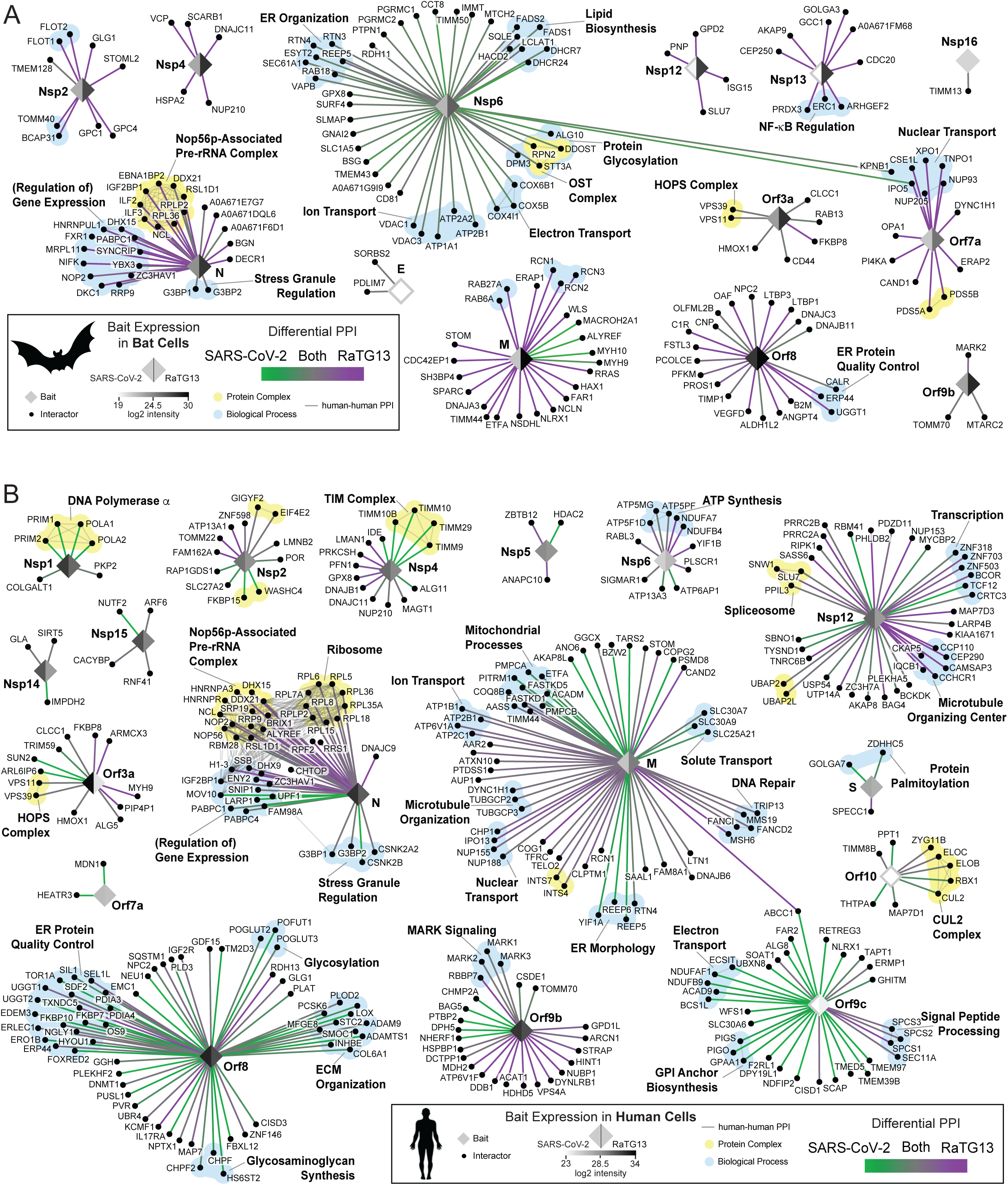
Conserved and distinct host-virus interaction landscapes of SARS-CoV-2 and RaTG13. (A) Comparative differential interaction map depicting SARS-CoV-2-RaTG13 comparison of bat-virus interactions. Edge color represents the differential interaction score (DIS): green and purple indicate interactions specific to SARS-CoV-2 and RaTG13, respectively, while grey indicates shared interactions. Each bait node is split into two halves, colored by the log₂ intensity values representing bait expression for SARS-CoV-2 (left) and RaTG13 (right). Thin grey edges indicate known host-host (human-human) physical interactions. Host proteins that belong to the same protein complex or biological process are highlighted in light yellow and light blue, respectively. Host-host interactions, complex annotations, and biological process groupings were derived from CORUM, Gene Ontology (biological process), and manual curation from the literature. (B) Comparative differential interaction map depicting virus-host interactions for SARS-CoV-2 and RaTG13 in human cells.

Some of the host- or virus-specific interactions might result from increased sensitivity of the immunoprecipitations due to differences in viral and host protein expression, or to peculiarities of the cell lines used. Nevertheless, these comparative interactome maps across bat and human cells delineate conserved host interactions exploited by sarbecoviruses. This dataset is likely to include virus-specific interactions that influence host tropism, immune evasion, and cross-species transmission. Thus, our data provide valuable information for further validation to identify host and viral factors involved in sarbecovirus spillovers from bats to humans.

### Differential interaction of Orf9b from SARS-CoV-2 and RaTG13 with mitochondrial proteins Tom70 and MTARC2

To further dissect virus- and host-specific differences, we examined the global distribution of DIS scores across all interactions (**Figure 3A, B**). In agreement with our earlier findings (**Figure S2F**), DIS values for cross-host species comparisons were enriched near ±1, indicating a higher degree of divergence for interactions across different hosts (**Figure 3A**, light and dark blue). In contrast, DIS values for comparisons between SARS-CoV-2 and RaTG13 within the same host were closer to zero, which indicates more shared interactions (**Figure 3A**, red and orange). This analysis revealed individual host proteins with strong virus-specific and/or host-specific enrichment. Notably, we identified a RaTG13-specific interaction of Orf7a with proteins involved in nuclear transport (i.e. CSE1L, NUP205, NUP93, TNPO1, and XPO1) in both bat and human cells (**Figure 3B**). Several bat-specific interactions were identified, which may provide insights into distinct immune modulation strategies or mechanisms of viral persistence in bat hosts (**Figure 3B, S4A**). Some of these bat-specific interactions involve ISG15, SCARB1, and PRDX3, which may reflect unique viral strategies for immune modulation^25–28^. To investigate if the absence of these interactions in HEK293T cells is due to lacking or low expression of the corresponding host proteins, we performed deep proteome profiling to compare host protein abundance levels across cell types. Interestingly, a substantial number of bat-specific host proteins (e.g. NIFK, PRDX3, SCARB1, BSG and MTARC2) were expressed at similar or even higher levels in HEK293T cells (**Figure S4A**), suggesting that differential expression was not the cause of lack of interactions; instead, factors such as sequence divergence, post-translational modifications, or cell-type-specific complexes are likely to explain these findings. When comparing all four data sets, we identified several conserved interactions, underscoring the critical involvement of these shared host proteins and complexes in sarbecovirus infection (**Figure 3B, S4B**). Notably, the N protein consistently interacted with stress granule-associated proteins, a previously characterized interaction known to disrupt stress granules (G3BP1, G3BP2) and suppress the interferon response^29^. We also detected a conserved interaction between the viral protein Orf3a and HOPS complex components VPS11 and VPS39, which have been shown to inhibit autophagy^30^. Additionally, we observed a conserved interaction of Orf9b with Tom70 (gene name TOMM70), a mitochondrial import receptor essential for MAVS signaling, which we previously identified as a direct target of SARS-CoV-2 Orf9b in human cells for suppression of innate immune responses^23,31^.

**Figure 3.**
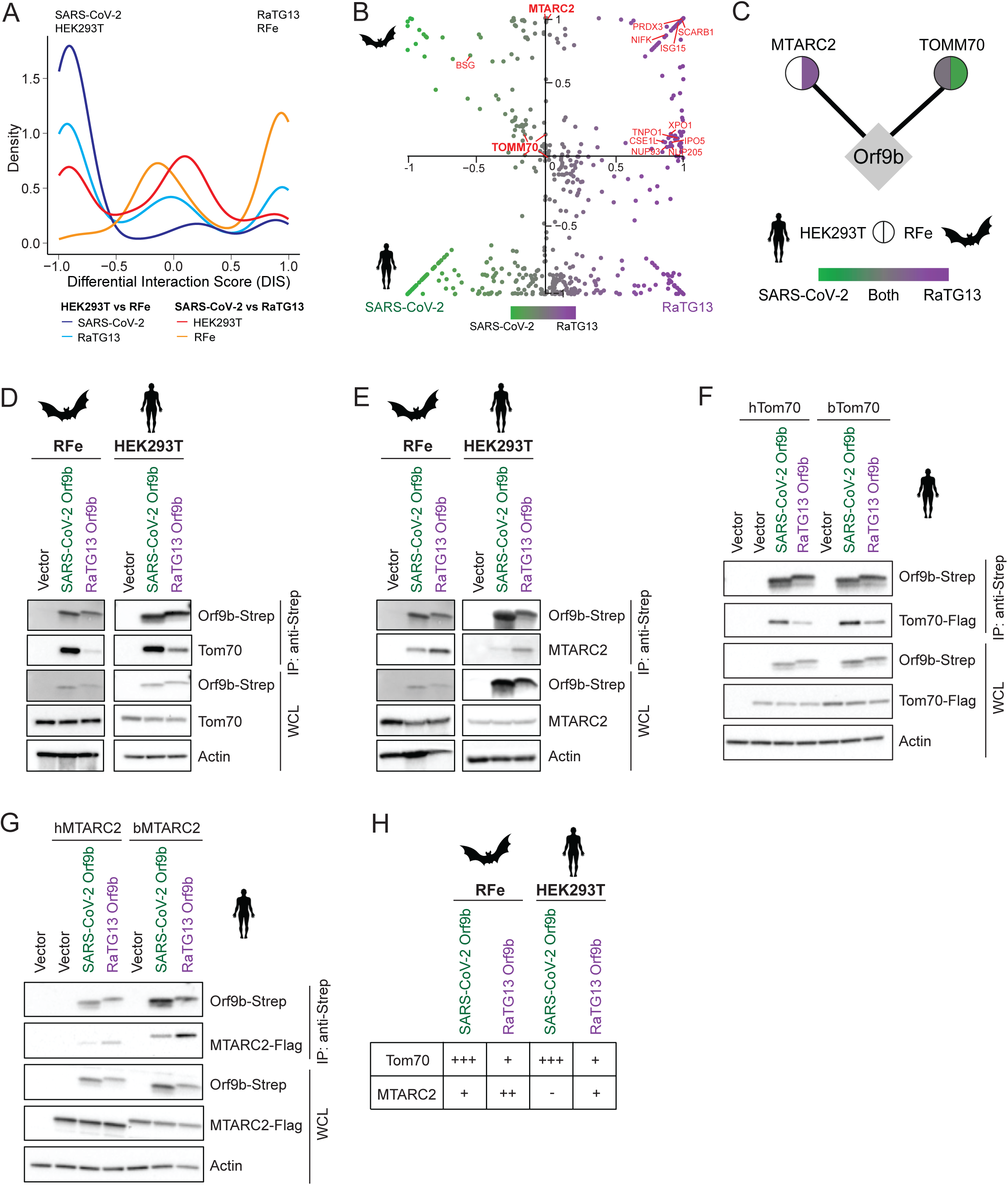
Differential interaction of SARS-CoV-2 and RaTG13 Orf9b with mitochondrial proteins Tom70 and MTARC2 across bats and human. (A) Density histogram representing the distribution of Differential Interaction Scores (DIS) comparing SARS- CoV-2 and RaTG13 interactomes within and across HEK293T and RFe cells. The PPIs of identical baits between SARS-CoV-2 and RaTG13 were excluded from the comparison across viruses. (B) Scatter plot of Differential Interaction Scores showing virus- and species-specific host protein interactions for SARS-CoV-2 and RaTG13 in bat (top) and human (bottom) cells. Each dot represents a host protein, colored by viral specificity (green for SARS-CoV-2, purple for RaTG13 and gray for shared). (C) Comparative interaction network of Orf9b from SARS-CoV-2 and RaTG13 with mitochondrial proteins Tom70 and MTARC2. Each node represents a host interactor, with split circles indicating enrichment in HEK293T (left half) and RFe cells (right half). Node color reflects the log₂FC of prey enrichment for RaTG13 relative to SARS-CoV-2, based on MSstats analysis: green for SARS-CoV-2-specific, purple for RaTG13- specific, gray for shared interactions, and white for interactions that were not detected. (D, E) Co-immunoprecipitation (co-IP) of Strep-tagged Orf9b from SARS-CoV-2 and RaTG13 with endogenous Tom70 (D) or MTARC2 (E) in HEK293T and RFe cells. HEK293T cells were transfected with Orf9b-Strep, and IP was performed using Strep-Tactin magnetic beads. For co-IP in RFe cells, doxycycline- inducible stable cell lines expressing Orf9b were used. Representative western blots of whole cell lysates (WCL) and IP eluates are shown. β-Actin was used as a loading control for WCLs. (F) Co-IP of Strep-tagged Orf9b with transiently expressed Flag-tagged Tom70 from human (hTom70) or bat (*Rhinolophus ferrumequinum*, bTom70) in HEK293T cells. Cells were transfected with Tom70-Flag alone or with Orf9b-Strep, as indicated. IP was performed using anti-Strep antibodies. (G) Co-IP of Strep-tagged Orf9b with Flag-tagged MTARC2 from human (hMTARC2) or bat (*Rhinolophus ferrumequinum*, bMTARC2) in HEK293T cells. Cells were transfected with MTARC2-Flag alone or with Orf9b-Strep, as indicated. (H) Summary table of interaction strength between Orf9b (SARS-CoV-2 and RaTG13) and Tom70 or MTARC2 from human and bat, based on co-IP results.

To uncover functional modules perturbed by each virus, we performed network propagation analysis on the PPIs to reveal proximal pathways and protein complexes that may not be directly captured by AP-MS alone. Due to lower bait expression and sparse PPI detection for SARS-CoV-2 in bat cells, we performed this analysis on the human datasets, comparing SARS-CoV-2 and RaTG13 PPIs to identify both virus-specific and shared pathway-level signatures. Using either virus as seed in the Reactome functional interactions (FI) network, we identified numerous virus-specific and shared proteins, including novel candidates that were not detected in the PPI dataset (**Figure S5A**). Propagation analyses revealed a prominent enrichment of mitochondrial-associated processes for SARS-CoV-2, including oxidative phosphorylation, ATP synthesis, mitochondrial electron transport, and targeting/localization to the mitochondrion, as determined by gene set over-representation analysis of propagated genes (**Figure S5B**). To pinpoint viral proteins potentially driving the observed mitochondrial pathway associations, we next looked at the number of viral baits whose interactors were linked to mitochondrial-related GO terms by performing network propagation separately for each viral protein. This analysis highlighted SARS-CoV-2 Orf9b and Nsp4 as top candidates, consistent with previous reports implicating SARS-CoV-2 Orf9b in the modulation of mitochondrial function and antiviral signaling^31,32^. In contrast, RaTG13 Orf9b exhibited fewer mitochondrial associations, suggesting potential functional divergence (**Figure S5C-E**). To further explore the interaction landscape of Orf9b, we conducted a separate quantitative and comparative PPI mapping of SARS-CoV-2 and RaTG13 Orf9b in both bat and human cells simultaneously (**Table S5**, see Methods section ‘Orf9b PPI’). This analysis revealed differential interactions of SARS-CoV-2 and RaTG13 Orf9b with the mitochondrial proteins Tom70 and MTARC2, particularly in bat cells (**Figure 3C**). Although the Orf9b-Tom70 interaction is conserved across both viruses in human cells, SARS-CoV-2 interacts with Tom70 more strongly than RaTG13 in bat cells. Conversely, we found that the MTARC2-Orf9b interaction was detected only in bat cells and was enriched for RaTG13 Orf9b over SARS-CoV-2 Orf9b. MTARC2 functions as a mitochondrial amidoxime-reducing component involved in metabolic processes such as lipid metabolism and redox regulation^33,34^. However, it has not previously been described to play a role in viral pathogenesis.

We next validated our proteomics findings using Strep-tagged co-immunoprecipitation (co-IP) assays and confirmed that SARS-CoV-2 Orf9b interacts with endogenous Tom70 in both bat and human cells, but RaTG13 Orf9b only weakly interacts with Tom70 in both cell-types (**Figure 3D**). We also confirmed the stronger interaction of RaTG13 Orf9b with endogenous MTARC2 in bat cells; this interaction is also consistently observed in human cells, albeit more weakly (**Figure 3E**).

Bat and human Tom70 and MTARC2 orthologs share 96.88% and 76.72% sequence identity, respectively, whereas Orf9b is 92.78% identical between SARS-CoV-2 and RaTG13, differing by seven amino acids. To directly compare binding strength of Orf9b across bat and human ortholog proteins, we performed co-IPs with Strep-tagged SARS-CoV-2 or RaTG13 Orf9b upon co-expression of Flag-tagged Tom70 proteins from *R. ferrumequinum* (bTom70) or humans (hTom70) in HEK293T cells. We found that SARS-CoV-2 Orf9b interacted equally well with both human and bat Tom70 orthologs, while RaTG13 Orf9b showed weaker binding to both orthologs (**Figure 3F**). Similarly, we conducted co-IPs with Strep-tagged SARS-CoV-2 or RaTG13 Orf9b upon co-expression of Flag-tagged MTARC2 from *R. ferrumequinum* (bMTARC2) and human (hMTARC2). In each case, MTARC2 co-precipitated more strongly with RaTG13 Orf9b compared to SARS-CoV-2 Orf9b (**Figure 3G**). Additionally, we observed a stronger interaction between RaTG13 Orf9b and bMTARC2 compared to hMTARC2, consistent with our previous results. In conclusion, we observed that Orf9b has switched preference in binding to Tom70 (SARS-CoV-2) versus MTARC2 (RaTG13) (summarized in **Figure 3H**), suggesting functional differences in mitochondrial targeting, immune evasion mechanisms, and host adaptation between both viruses.

### Orf9b residue 72 as a molecular switch governing its interactions with Tom70 and MTARC2

We and others previously resolved the structure of SARS-CoV-2 Orf9b in complex with the C-terminal domain of hTom70^23,35^. SARS-CoV-2 Orf9b alone forms a dimer and consists mainly of β-sheets [PDB: 6Z4U]^36^. However, upon binding to Tom70 in a 1:1 stoichiometry^35^, Orf9b adopts an α-helical conformation and interacts with the substrate-binding site of Tom70. This Orf9b-Tom70 complex is stabilized by hydrogen bonding and hydrophobic interactions. Using these experimentally determined structures as templates, we utilized homology modeling to predict the binding interface between RaTG13 Orf9b and hTom70 and compared key contact residues and binding properties. Previous structural studies suggested that SARS-CoV-2 Orf9b residue T72 is situated in a hydrophobic pocket of Tom70, engaging in extensive hydrophobic interactions^23^. Interestingly, SARS-CoV-2 Orf9b T72 forms a hydrogen bond with V556 in Tom70, whereas the corresponding I72 in RaTG13 Orf9b does not form any hydrogen bonds with Tom70 (**Figure 4A**). Instead, the neighboring RaTG13 Orf9b residue R70 forms a hydrogen bond with Tom70 residue G218; this interaction is absent in the SARS-CoV-2 Orf9b-Tom70 structure. Secondary structure predictions using JPred^37^ indicate that while both sequences retain an overall helical structure, the RaTG13 variant (R70/I72) exhibits an extended predicted α-helix compared to SARS-CoV-2 (Q70/T72), suggesting that these substitutions modestly enhance local helical propensity (**Figure S6A**). These observations may explain the differential binding properties of RaTG13 versus SARS-CoV-2 Orf9b with Tom70.

**Figure 4.**
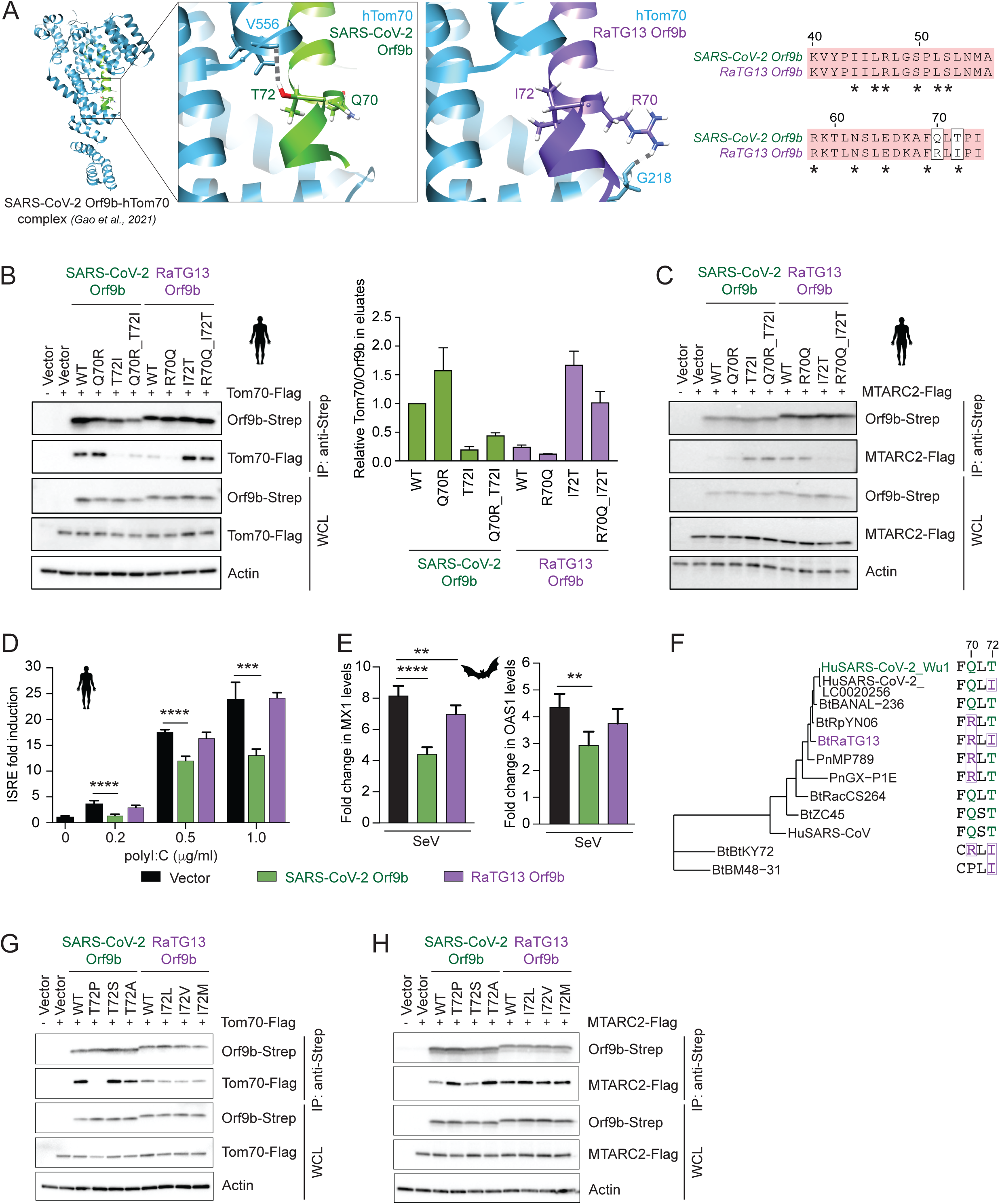
Molecular switch at residue 72 in Orf9b governs interactions with Tom70 and MTARC2 to modulate antiviral response. (A) Comparative structural modeling of RaTG13 Orf9b (purple) bound to human Tom70, using the crystal structure of SARS-CoV-2 Orf9b (green) in complex with Tom70 as a template (PDB IDs 7KDT and 7DHG). As per convention, hydrogen atoms are shown in white, oxygen in red and nitrogen in blue on selected side chains of both Orf9b structures. Hydrogen bonds are indicated by a grey broken line. The right panel shows a sequence alignment of SARS-CoV-2 and RaTG13 Orf9b. Conserved residues are shaded in red; asterisks indicate residues in SARS-CoV-2 Orf9b involved in interaction with Tom70. Residues at positions 70 and 72 (boxed) were selected for mutational analysis. (B) Co-immunoprecipitation (co-IP) of Strep-tagged Orf9b mutants with Flag-tagged Tom70 in HEK293T cells. Western blots (left) show Tom70 and Orf9b levels in eluates and whole cell lysates (WCL). SARS-CoV- 2 and RaTG13 mutants include single and double point mutations at positions 70 and 72. Bar graph (right) shows densitometric quantification of Tom70 co-precipitation normalized to Orf9b levels, averaged from biological replicates (mean ± SD). (C) Co-IP of Flag-tagged MTARC2 with SARS-CoV-2 and RaTG13 Orf9b wild-type and mutants in HEK293T cells. (D) ISRE reporter activity in HEK293T cells expressing Orf9b at varying poly I:C concentrations. Strep-tagged Orf9b from SARS-CoV-2 or RaTG13 was overexpressed in HEK293T cells containing a Lucia reporter under the control of the IFN-β/ISG56 promoter, alongside a vector control. Fold activation was calculated relative to unstimulated cells. ***P<0.001, ****P<0.0001. Statistical significance was calculated relative to vector control using ANOVA with Tukey’s multiple comparisons test. (E) Fold changes in Sendai virus (SeV)-induced ISG levels in bat cells upon Orf9b expression. Doxycycline- inducible RFe stable cell lines expressing Orf9b were infected with SeV. RNA was isolated and analyzed by qRT-PCR to measure the levels of MX1 and OAS1. RFe cells expressing empty vector were used as a control and fold changes were calculated relative to uninfected cells. The data represent the mean ± SD from one representative experiment (n=3). **P<0.01, ****P<0.0001. Statistical significance was calculated relative to vector control using ANOVA with Tukey’s multiple comparisons test. (F) Phylogenetic tree of 12 diverse coronavirus genomes, generated using a nucleotide alignment of the N open reading frame. A 4aa region of the Orf9b sequence of each virus is shown, with changes at residues 70 and 72 highlighted. The T72I mutation arose independently in RaTG13 and in an isolate of human SARS- CoV2 (“LC0020256”) (G, H) Co-IP of Flag-tagged human Tom70 (G) or MTARC2 (H) with SARS-CoV-2 and RaTG13 Orf9b wild- type and mutants in HEK293T cells.

To further explore these structural differences, we assessed the impact of specific residues on Orf9b interactions by generating point mutants of SARS-CoV-2 and RaTG13 Orf9b, and tested them in co-IP assays upon co-expression with Flag-tagged hTom70. While mutating residue Q70 in SARS-CoV-2 Orf9b (Q70R) did not change the interaction, mutation of T72 (T72I) strongly impaired its interaction with Tom70. Conversely, mutating RaTG13 Orf9b residue 72, both alone (I72T) or as a double mutant (R70Q/I72T), restored Tom70 binding to levels similar to SARS-CoV-2 Orf9b (**Figure 4B**). We also observed a modest increase in Tom70 binding with the Q70R mutation in SARS-CoV-2 Orf9b when compared in the context of the T72I background. Similarly, the R70Q mutation in RaTG13 Orf9b further reduced Tom70 interaction relative to wild-type (WT). These findings are consistent with our structural modeling data. We also examined interactions of these mutants with human MTARC2. In line with the above results, we found that the RaTG13 Orf9b-MTARC2 interaction was disrupted by the I72T mutation but was unaltered by the R70Q mutation (**Figure 4C**). Similarly, the T72I mutation in SARS-CoV-2 Orf9b conferred a gain of interaction with MTARC2. These findings suggest that amino acid residue at position 72 in Orf9b acts as a molecular switch, switching Orf9b interaction between Tom70 and MTARC2.

Given the role of SARS-CoV-2 Orf9b in suppressing the innate immune response through its interaction with Tom70^31,32^, we further assessed how RaTG13 Orf9b affects innate immune activation downstream of RNA sensing. We expressed Orf9b from SARS-CoV-2 or RaTG13, alongside a vector control, in HEK293T cells stably expressing a Lucia reporter under the control of the IFNβ/ISG56 promoter, and stimulated the cells with poly(I:C). Consistent with our previous findings^31^, expression of SARS-CoV-2 Orf9b significantly inhibited reporter activation compared to the vector control, whereas no significant difference was observed in RaTG13 Orf9b-expressing cells (**Figure 4D**). This pattern was observed consistently across varying poly(I:C) concentrations tested in the assay. Technical limitations prevented us from performing the same assay in RFe cells. Instead, we performed qPCR to assess the expression levels of key ISGs in RFe cells expressing SARS-CoV-2 or RaTG13 Orf9b upon infection with Sendai virus. Compared to the vector control, MX1 and OAS1 induction was significantly reduced in SARS-CoV-2 Orf9b-expressing cells, whereas no significant difference was observed in RaTG13 Orf9b-expressing cells (**Figure 4E**). These results correlate with our co-IP findings that RaTG13 Orf9b exhibits reduced affinity for Tom70, suggesting that this weaker interaction may impair its function as an innate immune antagonist.

To better understand the evolutionary dynamics of Orf9b, we analyzed coronavirus genome sequences from both human and bat-derived isolates. Alignment of Orf9b protein sequences suggests that a threonine at position 72 (T72) is the ancestral residue for SARSr-CoVs: threonine is conserved across SARS-CoV-2 (HuSARS-CoV-2_Wu1), several closely related pangolin (Pn) and bat (Bt) viruses, and the distinct 2003 human SARS-CoV (**Figure 4F**). Interestingly, the threonine-to-isoleucine substitution (T72I) observed in RaTG13 (BtRaTG13) also arose independently in multiple lineages, including more divergent clades (e.g., BtKY72, BtBM48-31) and SARS-CoV-2 isolates (e.g., HuSARS-CoV-2_LC0020256); it was detected at low frequency in the human population throughout the pandemic (**Figure S6B, C**), consistent with previous estimates of its near-neutral fitness^38^. We also observed sequence variation at position 70, where arginine (R) is present in RaTG13 and several other bat and pangolin viruses, while glutamine (Q) is found in both SARS-CoV and SARS-CoV-2, as well as in their closest relatives. This observed pattern suggests that R70 was likely present in the common ancestor of RaTG13 and SARS-CoV-2, followed by a reversion to Q in the SARS-CoV-2 lineage.

Together, these findings suggest that Orf9b position 72 is evolutionarily constrained, with T72 broadly conserved across SARS-related viruses. To further investigate the functional significance of this residue, we introduced all possible single-nucleotide substitutions at codon 72 of SARS-CoV-2 or RaTG13 Orf9b, respectively, and assessed their impact on host protein interactions using co-IP assays (**Figure 4G**). Among these, T72P was the only additional SARS-CoV-2 mutant that showed a marked loss of interaction with Tom70, likely due to disruption of local protein structure. In contrast, T72A and T72S had minimal effects on Tom70 binding. Interestingly, we observed an opposite binding pattern with MTARC2, where the T72P and T72A mutant showed increased interaction comparable to SARS-CoV-2 WT Orf9b, while the T72S substitution showed similar binding (**Figure 4H**). These results suggest that residue 72 plays a differential role in modulating Orf9b interactions with distinct host targets, potentially through localized structural changes that selectively alter binding interfaces. Notably, only the T72I mutation is synonymous with respect to the N open reading frame, whereas all other nucleotide substitutions result in non-synonymous changes in N. This dual coding constraint may further limit acceptable mutations at this position. Together with the low frequency of this mutation in the human population, our results suggest that T72I might be a deleterious mutation, at least in human cells. Altogether, these findings highlight the dual genomic and functional constraints shaping Orf9b evolution, arising from its role in host interaction and its genomic overlap with the N coding sequence.

### A single nucleocapsid substitution supports productive SARS-CoV-2 Infection in RFe-AT cells

We next sought to investigate the role of Tom70 and MTARC2 during infection in bat cells. However, immortalized RFe cells were not permissive to SARS-CoV-2 infection under multiple experimental conditions. To overcome this block, we generated a RFe cell line that stably expressed human ACE2 and TMPRSS2 (which we named RFe-AT), and confirmed expression of both human proteins by Western blot (**Figure 5A**). To determine whether viral entry is blocked in parental RFe cells, we produced HIV-1-based pseudotyped particles expressing a luciferase reporter and a SARS-CoV-2 WA1 spike protein lacking the 19 amino acids at the C-terminus of the cytoplasmic tail, important for the ER/Golgi retention (WA1 SΔ19)^39^. Entry assay results showed that overexpression of hACE2 and hTMPRSS2 was sufficient to overcome the entry restriction in RFe cells. As expected, control virus-like particles pseudotyped with the VSV-G protein efficiently infected both RFe and HEK293T cells, independent of hACE2 and hTMPRSS2 expression (**Figure 5B**). These results suggest that SARS-CoV-2 spike has reduced binding affinity for endogenous RFe ACE2, or that RFe cells express very low levels of endogenous ACE2, consistent with reports from other bat cell lines^40^, thereby limiting viral entry. Notably, entry of both HIV-1 based VLPs was reduced in RFe cells compared to HEK293T cells, suggesting the presence of additional barriers to viral entry in RFe cells that warrant future investigation. To assess whether limited RFe ACE2 expression contributes to this restriction, we overexpressed HA-tagged ACE2 from *R. ferrumequinum* (bACE2) or human (hACE2) in HEK293T cells, which are not susceptible but are permissive to SARS-CoV-2 infection, and measured the percentage of nucleocapsid protein (N) positive cells within the ACE2 positive population at 24 hours post-infection with SARS-CoV-2 WA1 (**Figure 5C**). As expected, expression of hACE2 supported efficient SARS-CoV-2 entry and replication, whereas even overexpression of bACE2 could not overcome the entry block. These results, together with previously published data^41^, suggest that bACE2 is unable to mediate SARS-CoV-2 entry, likely due to the low amino acid identity of key residues required for interaction with the spike RBD.

**Figure 5.**
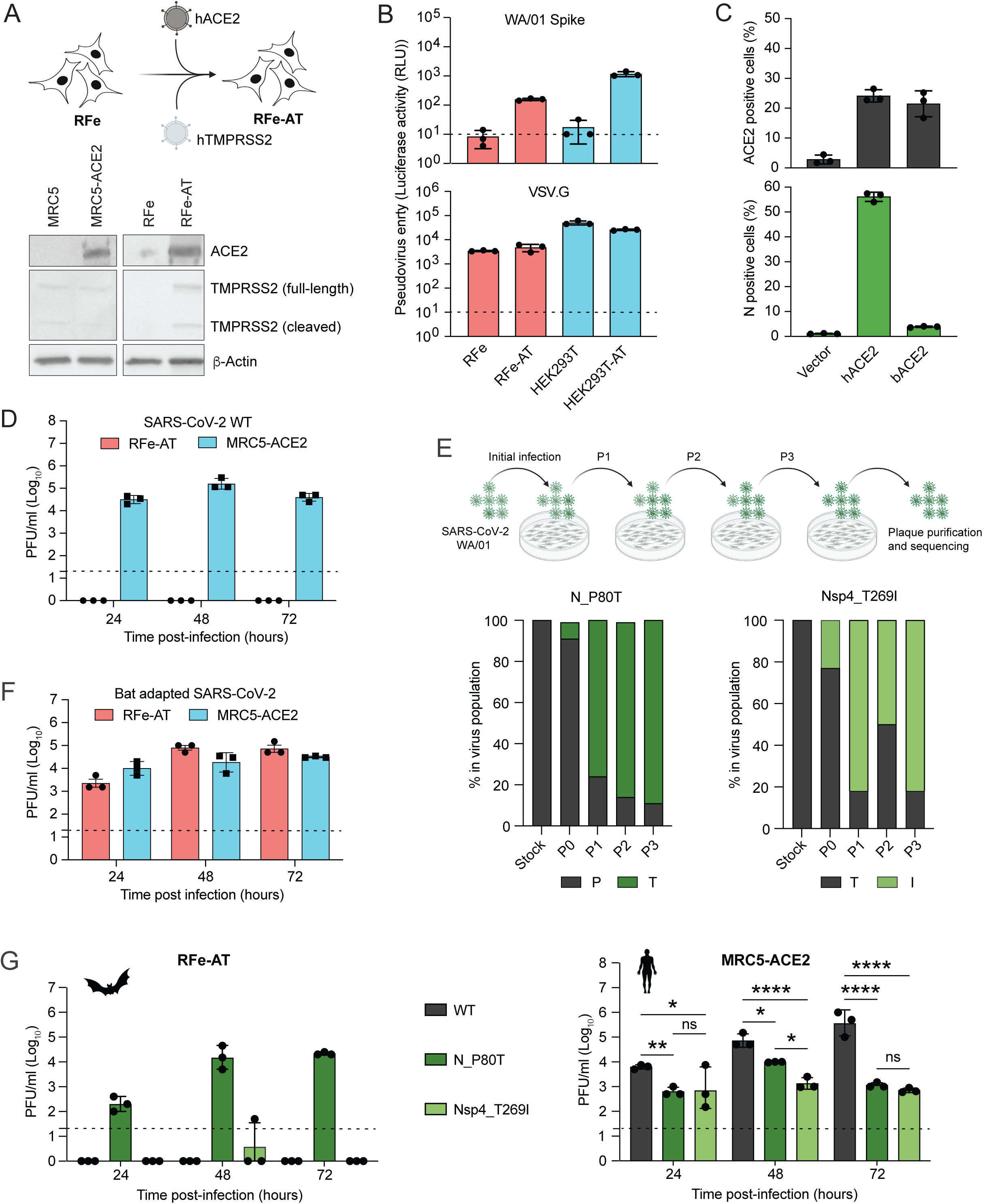
Establishment of a Bat Cell Infection Model through hACE2/hTMPRSS2 Expression and SARS-CoV-2 Adaptation. (A) Schematic illustrating the generation of the RFe-AT cell line by introducing human ACE2 and TMPRSS2 into immortalized RFe cells (top). Western blot analysis (bottom) shows expression of human ACE2 and TMPRSS2 (full-length and cleaved forms) in the indicated cell lines. β-actin serves as a loading control. (B) Functional assessment of SARS-CoV-2 entry using pseudotyped HIV based particles. Luciferase-based entry assays were performed using HIV-based particles pseudotyped with SARS-CoV-2 WA/01 Spike (top) or VSV-G (bottom) in RFe, RFe-AT, HEK293T, and HEK293T-AT cells. Firefly luciferase activity is shown as relative luminescence units (RLU). (C) Assessment of SARS-CoV-2 entry mediated by ACE2 orthologs from human (hACE2) or bat (bACE2). HEK293T cells were transfected with vector control, hACE2, or bACE2, followed by infection with SARS- CoV-2 WA/01 at MOI 1. Top panel depicts the percentage of ACE2-positive cells determined by flow cytometry. Percentage of infected (N-positive) cells in ACE2-positive population was measured by intracellular staining for viral nucleocapsid (N) protein (bottom). Data represent ± SD from one of three independent experiments. (D) Replication kinetics of SARS-CoV-2 WA/01 in RFe-AT (red) and MRC5-hACE2 (blue) cell lines. Cells were infected with the virus at MOI 0.1. Virus supernatant was collected at 24, 48 and 72 h post-infection and plaque assays were performed in Vero E6 cells. (E) Schematic overview of the SARS-CoV-2 adaptation experiment in bat cells (top). The virus was serially passaged (P1-P3), followed by plaque purification and viral genome sequencing. Bar plots (bottom) show the frequency of specific missense mutations (N_P80T and Nsp4_T269I) in the viral population across passages. (F) Replication kinetics of RFe adapted SARS-CoV-2 WA1 virus in RFe-AT and MRC5-hACE2 cell lines. Cells were infected at MOI 0.1 in triplicates. Supernatant was collected at 24, 48 and 72 h post-infection and plaque assays were performed in Vero E6 cells. (G) Growth kinetics of recombinant SARS-CoV-2 wild-type (WT) and viruses carrying mutations leading to N_P80T or Nsp4_T269I amino acid substitutions in RFe-AT (left) and MRC5-hACE2 (right) cells. Cells were infected at MOI 0.1, and virus titers in supernatants were measured by plaque assay at the indicated time points. Data are presented as mean ± SD from one representative biological replicate (n = 3). *P < 0.05, **P < 0.005, ****P < 0.00005. Statistical significance was determined using one-way ANOVA followed by Tukey’s multiple comparisons test.

To further characterize the susceptibility and permissiveness of the RFe-AT cell line to SARS-CoV-2, we performed infections at a MOI of 0.1 and measured viral growth kinetics over a 72-hour time course using plaque assays. Notably, we didn’t detect any productive infection in RFe-AT cells under these conditions. In contrast, robust and reproducible viral replication was observed at all time points in a human lung fibroblast cell line stably expressing human ACE2 (MRC5-ACE2), which was used as positive control (**Figure 5D**). These results indicate that, under these conditions, RFe-AT cells are not permissive to productive SARS-CoV-2 infection. To assess whether viral adaptation might overcome this restriction, we designed a high-MOI infection and serial passaging strategy. Specifically, we infected RFe-AT cells with SARS-CoV-2 WA/01 at an MOI of 1 and collected supernatant at 96 hours post-infection, which was used for three blind serial passages to facilitate potential adaptation to the bat cell environment. Such passaging could lead to the emergence of genetic changes (missense mutations, deletions, or insertions) that could specifically improve infection of RFe-AT cells. To identify such mutations, we isolated viral RNA from the supernatants of each passage and performed whole-genome sequencing of the SARS-CoV-2 viruses (**Figure 5E**). Strikingly, we identified three non-synonymous point mutations in the nucleocapsid (N_P80T), non-structural protein 4 (Nsp4_T269I), and spike protein (S_R682W; not shown) genes. The N_P80T nonsynonymous mutation also results in a synonymous mutation in Orf9b. The plaque-purified, bat-adapted virus carrying all three mutations enabled efficient replication in RFe-AT cells, comparable to that observed in MRC5-ACE2 cells (**Figure 5F**). Since all three mutations were previously found in SARS-CoV-2 circulating human isolates with low frequency^42^ and the S_R682W amino acid substitution has been previously reported as a common tissue culture adaptation^43^, we investigated the contribution of the other two substitutions to viral replication in RFe- AT cells. Using a previously established reverse genetics system^44^, we generated recombinant SARS-CoV- 2 WA/01 viruses carrying either the N_P80T or Nsp4_T269I amino acid substitution. Infection with these recombinant viruses revealed that the N_P80T mutation alone was necessary and sufficient to support productive replication in RFe-AT cells (**Figure 5G**). Moreover, replication of both recombinant mutants was significantly attenuated in MRC5-ACE2 cells (human) compared to wild-type SARS-CoV-2 WA/01, suggesting that these mutations confer host-specific adaptation to replication in bat cells while compromising replication efficiency in human cells.

### Loss of Tom70 Binding by Orf9b enhances innate immune activation

We applied our infection model to investigate the impact of Orf9b-Tom70/MTARC2 interactions on virus replication in the RFe-AT bat cell line. We rescued a bat adapted SARS-CoV-2 N_P80T virus carrying the T72I mutation in Orf9b (rSARS-CoV-2 N_P80T, Orf9b_T72I) and compared its growth kinetics to the rSARS-CoV-2 N_P80T and wild-type (WT) viruses in both bat and human cell lines. As expected, introduction of the Orf9b T72I mutation, which reduces the interaction between Orf9b and Tom70, led to reduced viral titers at 48 h post-infection in both RFe-AT and MRC5-ACE2 cell lines, as measured by plaque assays (**Figure 6A**). To further examine host innate immune responses, we performed bulk RNA-seq on RFe-AT and MRC5- ACE2 cells infected with WT and mutant viruses at 24 and 48 hours post-infection (**Figure 6B**). Transcriptome analysis revealed that, in MRC5-ACE2 cells, the Orf9b T72I mutant induced significantly stronger upregulation of interferon-stimulated genes (ISGs) at 24 hours post-infection compared to both WT and N_P80T viruses. This is consistent with our previous observations and other studies^31,32,45^ showing that Tom70 facilitates Orf9b-mediated suppression of innate immunity. The ISG response was much less pronounced in RFe-AT cells, suggesting either a dampened innate response in the bat cell line or delayed kinetics of immune activation. Overall, we also noticed lower viral replication in bat cells compared to human cells under the same MOI infection conditions, which could contribute to reduced induction of ISGs in bat cells. ISG expression profiling highlighted differential responses driven by mutant viruses across bat and human cell types (**Figure 6C, Table S6**). In RFe-AT cells, ISGs such as RSAD2 and DDX58 were modestly upregulated in response to infection, with relatively small differences between N_P80T and N_P80T, Orf9b_T72I. In contrast, MRC5-hACE2 cells showed robust induction of a broader panel of ISGs, particularly in response to the Orf9b_T72I virus, indicating reduced suppression of host antiviral responses due to the impaired interaction between Orf9b and Tom70.

**Figure 6.**
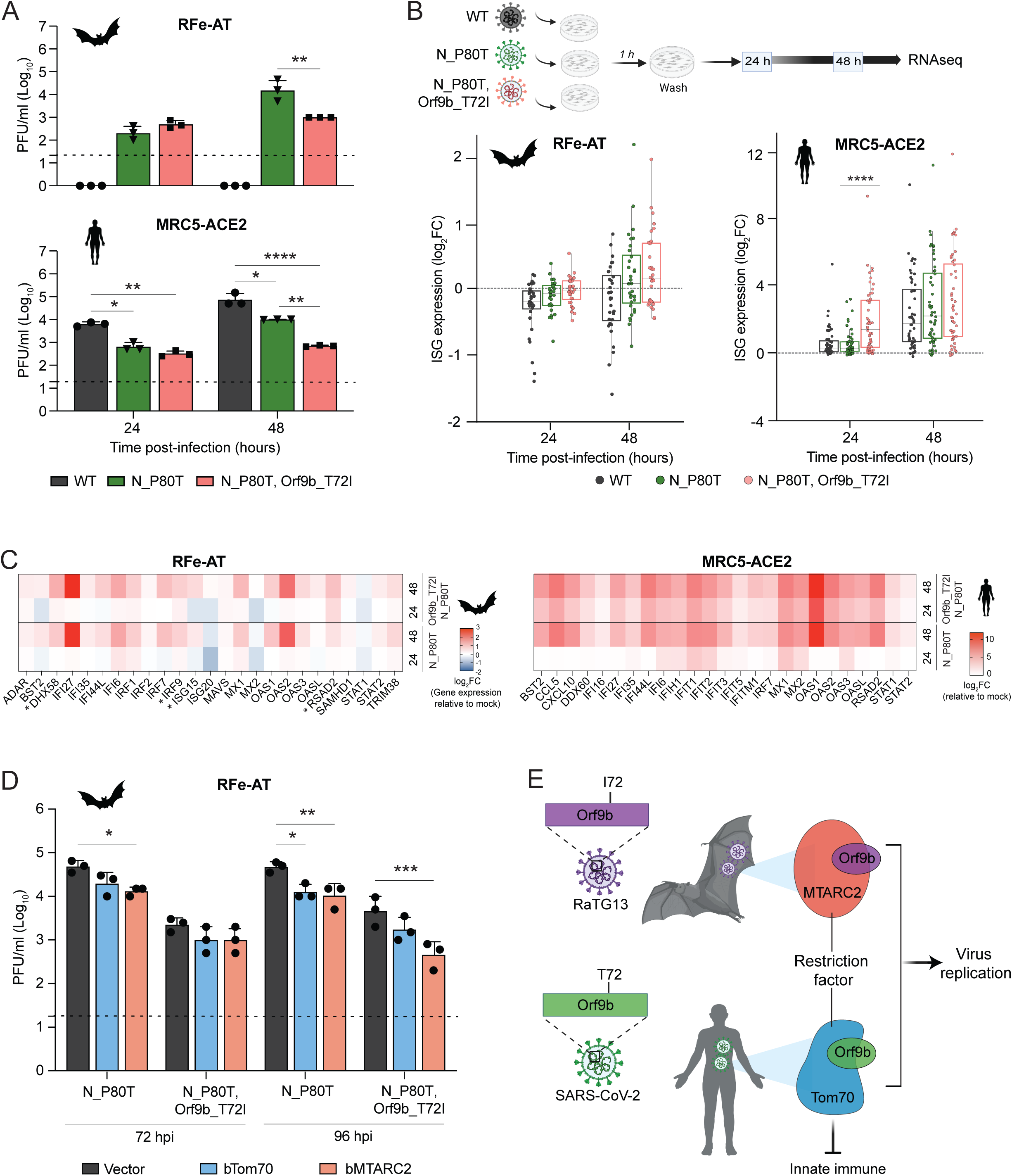
Orf9b T72I mutant lacking Tom70 binding modulates viral replication and immune signaling. (A) Bar graphs showing viral titers at 24 and 48 hours post-infection in RFe-AT or MRC5-ACE2 cells infected with the indicated viruses. Viral titers were measured by plaque assay on Vero E6 cells. Data are presented as mean ± SD from one representative experiment (n = 3). *P < 0.05, **P < 0.005, ****P < 0.00005. Statistical significance was determined using one-way ANOVA followed by Tukey’s multiple comparisons test. (B) Schematic of the experimental design (top): RFe-AT and MRC5-ACE2 cells were infected with the indicated viruses and harvested at 24 and 48 hours post-infection (hpi) for bulk mRNA sequencing. Box plots (bottom) show log₂-transformed fold change (log₂FC) of ISGs relative to uninfected controls at each time point. Each dot represents an individual ISG; boxes indicate the median (center line) and interquartile range (upper and lower bounds). (C) Heatmap displaying the log₂FC of expression levels of selected ISGs (from RNA-seq) in RFe-AT and MRC5-ACE2 cells infected with SARS-CoV-2 mutant viruses containing the N P80T mutation alone or in combination with the Orf9b T72I mutation. (D) Bar graphs showing viral titers at 72 and 96 h post-infection in RFe-AT cells stably expressing bTom70, bMTARC2, or vector control, following infection with the indicated viruses. Data represent mean ± SD from one representative experiment (n = 3). Statistical significance was assessed using one-way ANOVA with Tukey’s multiple comparisons test. *P < 0.05, **P < 0.01, ***P < 0.001. (E) Model illustrating divergent host interactions of Orf9b from RaTG13 and SARS-CoV-2. RaTG13 Orf9b, which contains isoleucine at position 72 (I72), interacts with the mitochondrial enzyme MTARC2 in bat cells and restricts virus replication. In contrast, SARS-CoV-2 Orf9b contains a threonine at the same position (T72) and binds to the mitochondrial import receptor Tom70 in human cells, leading to suppression of innate immune signaling. Our findings highlight species-specific adaptations of Orf9b that influence host immune modulation and viral replication.

To further investigate the functional impact of Tom70 and MTARC2 on SARS-CoV-2 replication in bat cells, we generated RFe-AT cell lines stably overexpressing either bat MTARC2 or Tom70, along with an empty vector control. Upon infection with the SARS-CoV-2 N_P80T virus, cell lines overexpressing either Tom70 or MTARC2 showed a modest reduction in viral titers compared to control cell lines (**Figure 6D**). These findings indicate that overexpression of either MTARC2 or Tom70 can enhance antiviral signaling or restrict viral replication to some extent. Notably, a more pronounced viral restriction was observed in MTARC2- expressing cells infected with the N_P80T, Orf9b_T72I mutant virus, which shows enhanced binding to MTARC2. These findings suggest a role for the MTARC2-Orf9b interaction in restricting viral replication in bat cells. In contrast, Tom70 overexpression did not result in a significant difference in viral titers compared to control following infection with the N_P80T, Orf9b_T72I mutant virus, as expected.

Our findings confirm previous studies, which showed that SARS-CoV-2 Orf9b leverages its interaction with the mitochondrial protein Tom70 to suppress host innate immune responses in human cells, thereby promoting viral replication (**Figure 6E**). In contrast, Orf9b from the bat coronavirus RaTG13, which contains an isoleucine at position 72 (I72), preferentially interacts with a different mitochondrial protein, MTARC2, in bat cells, where this interaction appears to restrict viral replication. Introducing the T72I mutation in SARS-CoV-2 Orf9b disrupts Tom70 binding while enhancing interaction with MTARC2, leading to impaired immune evasion and attenuated replication in both bat and human cell contexts. These results highlight a potential evolutionary trade-off in viral adaptation, where Orf9b fine-tunes its host interactions to switch binding between Tom70 in human cells and MTARC2 in bat cells to regulate replication and immune suppression in a species-specific manner.

## Discussion

The frequency of zoonotic spillovers of bat-borne viruses that cause severe disease in humans^46^ and other animals^47^ underscores the urgent need for bat-derived cellular models to investigate host-virus dynamics. These models will help to better understand the mechanisms that enable bats to serve as efficient viral reservoirs, ultimately informing future therapeutic strategies. In this study, we describe the generation of an immortalized lung fibroblast cell line derived from the Greater horseshoe bat (*Rhinolophus ferrumequinum*), a species within the *Rhinolophus* genus known to harbor SARS-like coronaviruses with pandemic potential.

To further study the relevance of our findings on virus infection in both human and bat cells, we created two additional resources. First, we rendered our bat RFe cell line susceptible to SARS-CoV-2 infection by overexpression of human ACE2 and TMPRSS2. Human ACE2 was essential to enable viral entry, consistent with previous findings that SARS-CoV-2 cannot use RFe ACE2 for entry^41^, nor ACE2 orthologs from other bat species^14^. However, no spread of infection was observed, indicating that viral adaptation is required to support efficient replication in bat cells. These findings suggest the presence of multiple host-specific barriers beyond receptor incompatibility that may restrict SARS-CoV-2 replication in bat cells. In this study, we identified a single mutation in the viral nucleocapsid sequence (N_P80T) that is sufficient to enable efficient replication of SARS-CoV-2 in *R. ferrumequinum* cells. While this mutation occurs within the region of the N gene that overlaps with the Orf9b gene, the nucleotide substitution is synonymous for Orf9b residue V76. Although the precise mechanism remains to be elucidated, this mutation may influence nucleocapsid-host interactions rather than intrinsic within-virus functions such as RNA binding or phase separation, which would generally be expected to have similar effects across hosts. For instance, N_P80T may modulate interactions with bat-specific immune sensors or stress granule components—processes known to impact viral replication and innate immune evasion^29,48^. SARSr-CoVs isolated from *Rhinolophus* bats, including RaTG13 used here, differ in several amino acids at the N protein near position 80. It might be possible that polymorphisms at the amino terminal RNA binding domain of the N protein are responsible for specific adaptation to different mammalian species, including bats. Ongoing work to characterize the broader set of viral mutations required for replication in *R. ferrumequinum* cells, along with the underlying mechanisms, is likely to reveal key species-specific restriction factors and inform risk assessments for the emergence potential of other bat-derived sarbecoviruses.

Through proteomics-based comparative interactome mapping in human and bat cells, we identified both shared and distinct protein-protein interactions for SARS-CoV-2 and its close bat coronavirus relative, RaTG13. We found that these viruses commonly target core cellular processes such as RNA metabolism, protein trafficking, and innate immune regulation. One of the conserved interactions between the viral protein Orf3a and HOPS complex components have been shown to inhibit autophagy^30^, consistent with a previous report that RaTG13 Orf3a interferes with human autophagy to similar levels as its SARS-CoV-2 counterpart^49^. Our study also revealed virus- and host-specific interaction patterns, which illustrate how subtle variations in viral proteins can lead to distinct molecular interactions and host response mechanisms. We identified a specific interaction between RaTG13 Nsp12 and ISG15 in bat cells. Recent work has shown that bat ISG15 harbors key residue changes across *Rhinolophus* species and exhibits stronger anti-SARS-CoV-2 activity compared to human ISG15^50^. Notably, despite the high sequence similarity between SARS-CoV-2 and RaTG13, we observed limited overlap in their interactomes within the same host species, suggesting functional divergence even among closely related sarbecoviruses. Understanding this divergence is essential for identifying species-specific vulnerabilities and informing the development of targeted therapeutic strategies.

Finally, we described how subtle differences in accessory protein Orf9b can reshape virus-host interactions across species. Comparative PPI and functional analyses demonstrated that SARS-CoV-2 Orf9b targets Tom70 to suppress innate immunity across host species. This is consistent with prior findings of this interaction’s role in mitochondrial immune modulation^23,31^, supporting a shared mechanism of immune suppression across species. In contrast, RaTG13 Orf9b preferentially interacts with MTARC2 in bat cells. MTARC2 regulates mitochondrial redox balance and may modulate antiviral signaling pathways by influencing ROS levels, inflammation, and interferon responses^34,51,52^. This interaction, leading to an MTARC2-mediated reduction in viral replication, may reflect Orf9b’s adaptation to the bat immune landscape, which is characterized by viral tolerance and reduced immunopathology. The functional relevance of these interactions is highlighted by our finding that a single amino acid substitution (T72I) in Orf9b disrupts Tom70 binding while enhancing MTARC2 interaction. When introduced into the bat-adapted virus, the MTARC2- favoring Orf9b mutation T72I resulted in impaired viral replication and heightened innate immune responses in both bat and human cells. Further, overexpression of MTARC2 resulted in reduced virus replication in bat cells, suggesting that MTARC2 acts as a host-specific restriction factor in the bats. Although MTARC2 has no known antiviral role in humans, its specific interaction with RaTG13 in bats points to a broader, yet uncharacterized, set of mitochondrial or metabolic regulators that may influence virus replication in bat reservoirs.

Interaction with MTARC2 is a derived trait observed in RaTG13 and a few other sarbecoviruses. Since this MTARC2 interaction was gained at the expense of the ancestral Tom70 interaction, it illustrates the importance of MTARC2 as a novel restriction factor, at least in bat cells. Interestingly, T72I is a synonymous mutation in the overlapping N ORF, whereas all other single-nucleotide substitutions at site 72 result in non-synonymous changes. Despite neutrality in the N ORF, T72I remains rare in human SARS-CoV-2 isolates, conceivably due to its disruption of Tom70 binding and impaired immune suppression. This highlights a dual evolutionary constraint at Orf9b position 72 imposed by overlapping coding sequences and the need to preserve Orf9b’s immune modulatory function. Together, these findings underscore the importance of viral accessory protein and host protein interactions in shaping viral fitness, immune modulation, and host adaptation, similar to poxviruses in which variations in accessory protein repertoires are related to host range and virulence^53^.

More broadly, our study highlights the utility of systematic host-virus interaction mapping across species as a powerful approach for dissecting the molecular underpinnings of host adaptation, immune evasion, and zoonotic emergence. Our comparative interactome maps serve as a resource for identifying both conserved cellular targets and species-specific interactors that may define host range barriers. Our findings underscore the evolutionary flexibility of accessory proteins like Orf9b in fine-tuning species-specific host interactions. The functional divergence between Orf9b orthologs in their ability to suppress host defenses may shape viral fitness and spillover potential. Future studies on defining the mechanistic role of newly identified bat-specific interactions, such as those involving MTARC2 and flotillins, and structurally characterizing these interfaces will provide insight into the evolutionary constraints governing these interactions. Systematic, cross-species interactome mapping – combined with functional infection models in reservoir hosts – offers a powerful framework for identifying molecular signatures predictive of zoonotic emergence. Expanding these approaches across diverse bat species and viral families will be essential for future pandemic preparedness.

## STAR Methods

### Key Resources Table

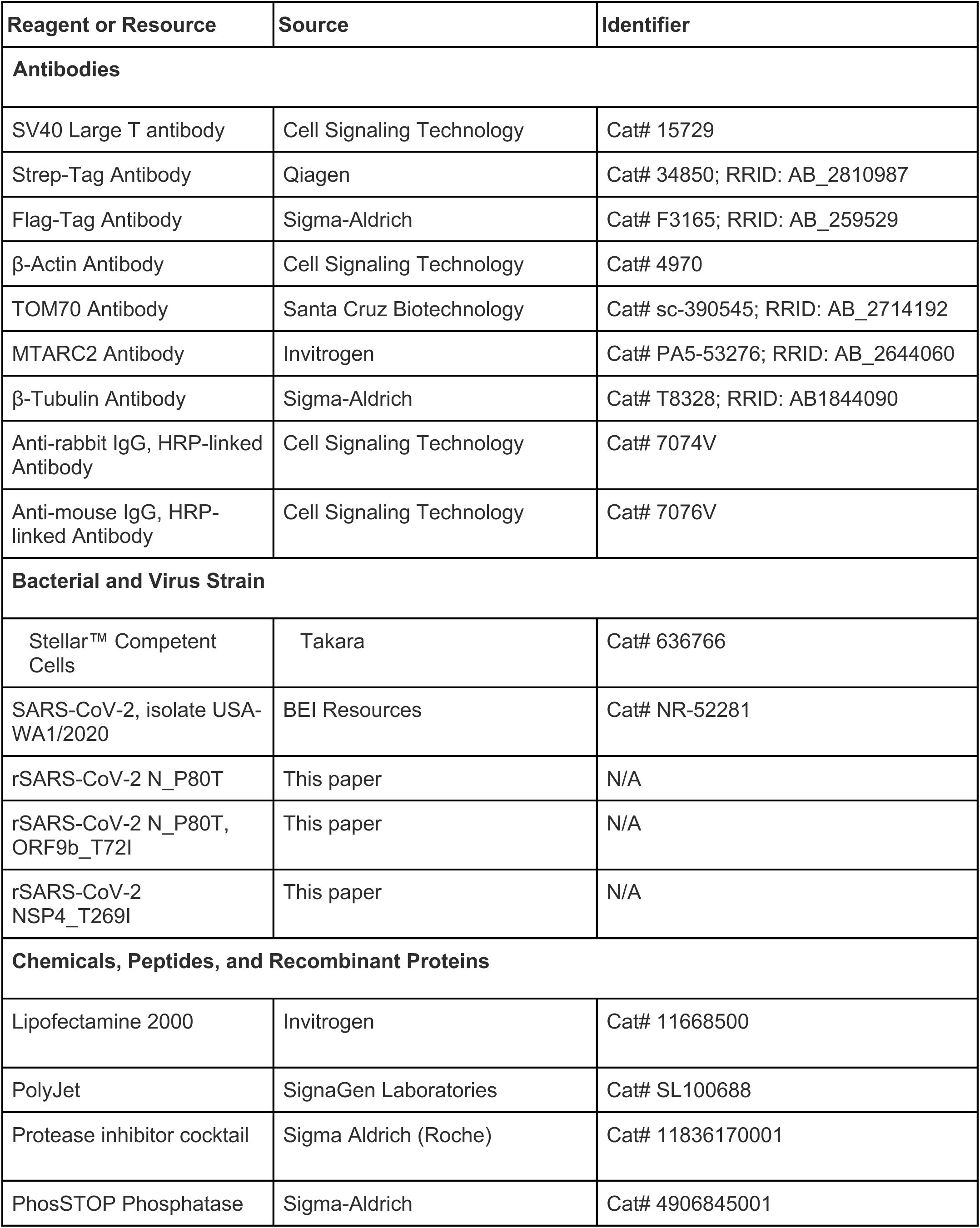
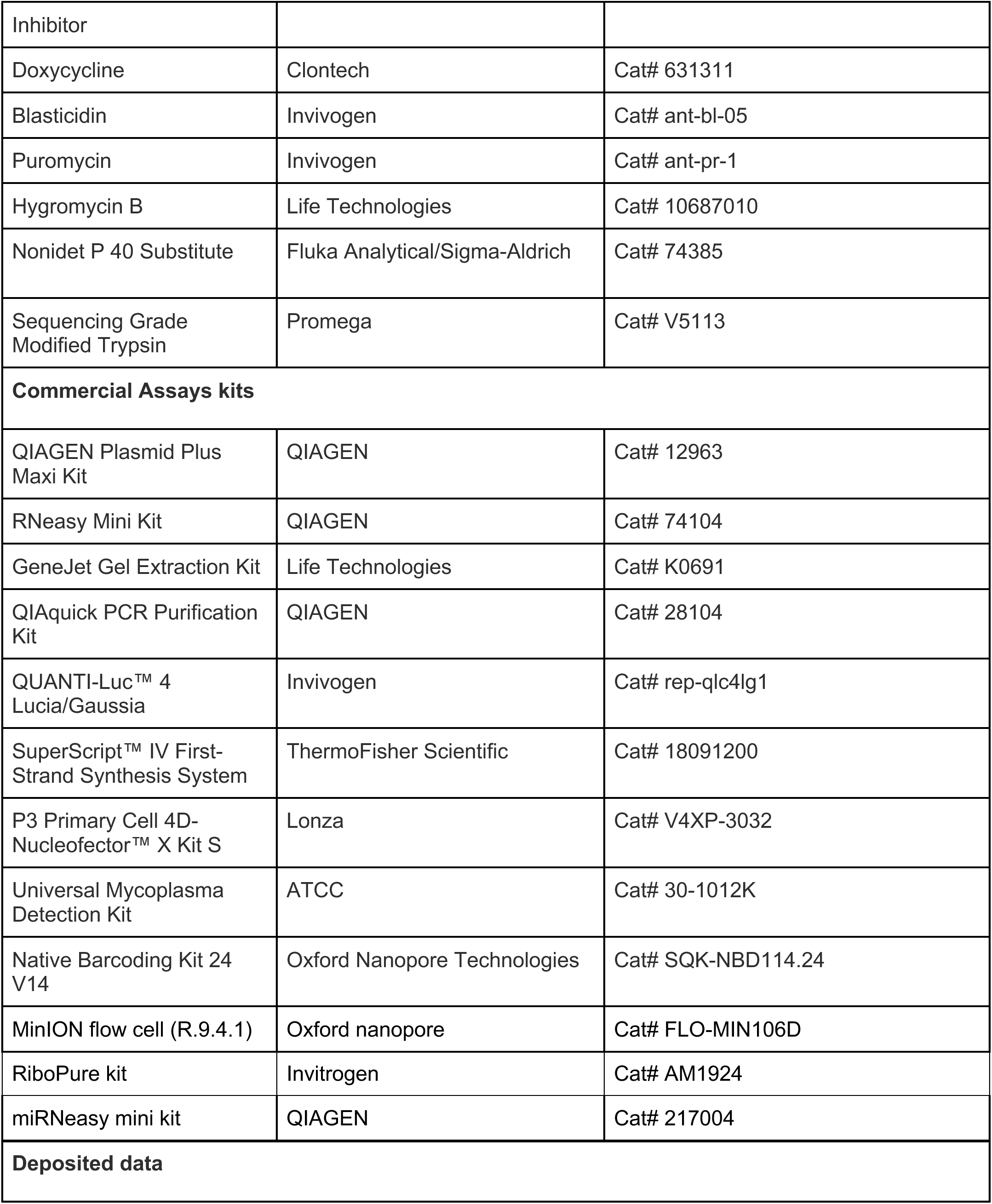
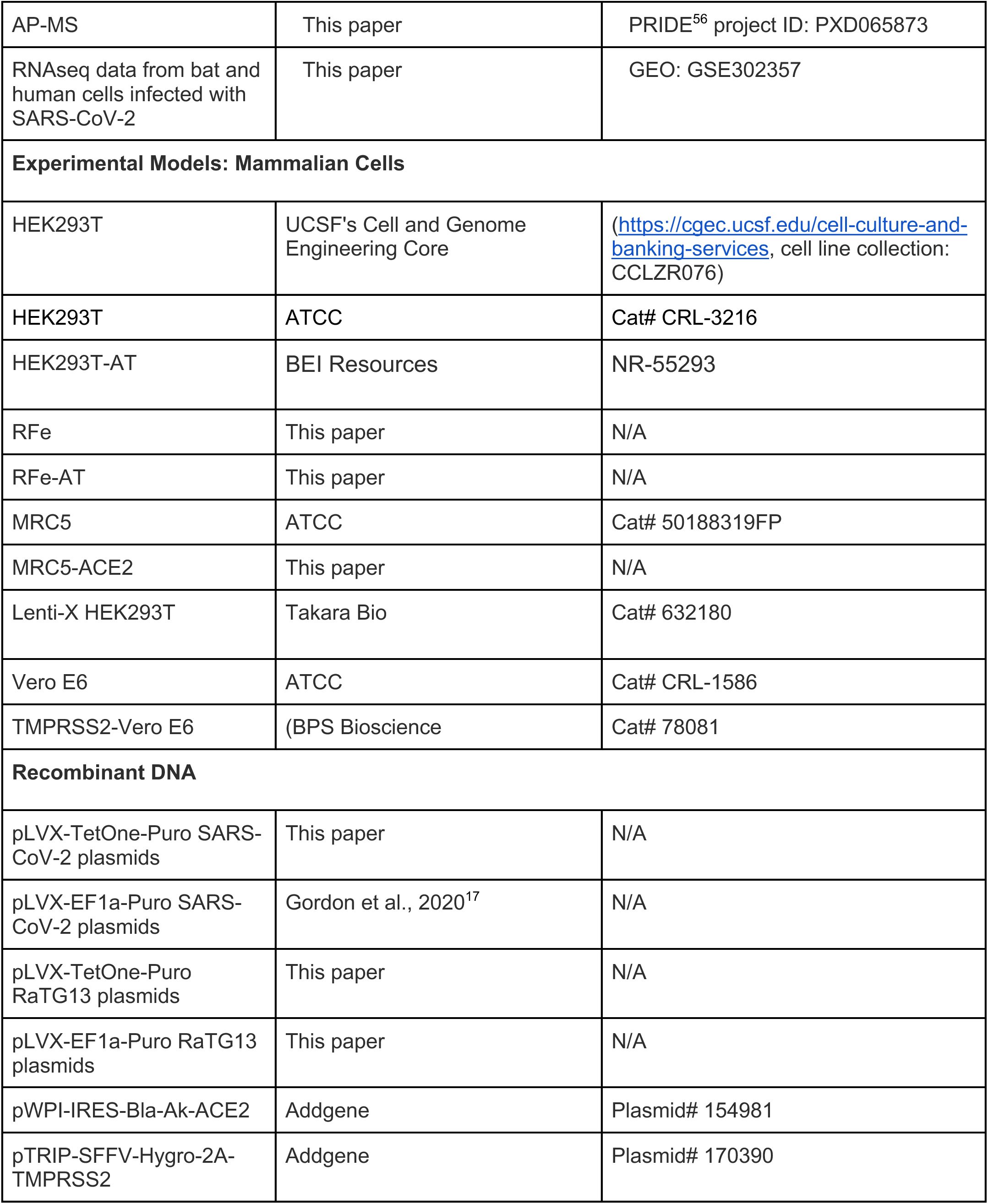
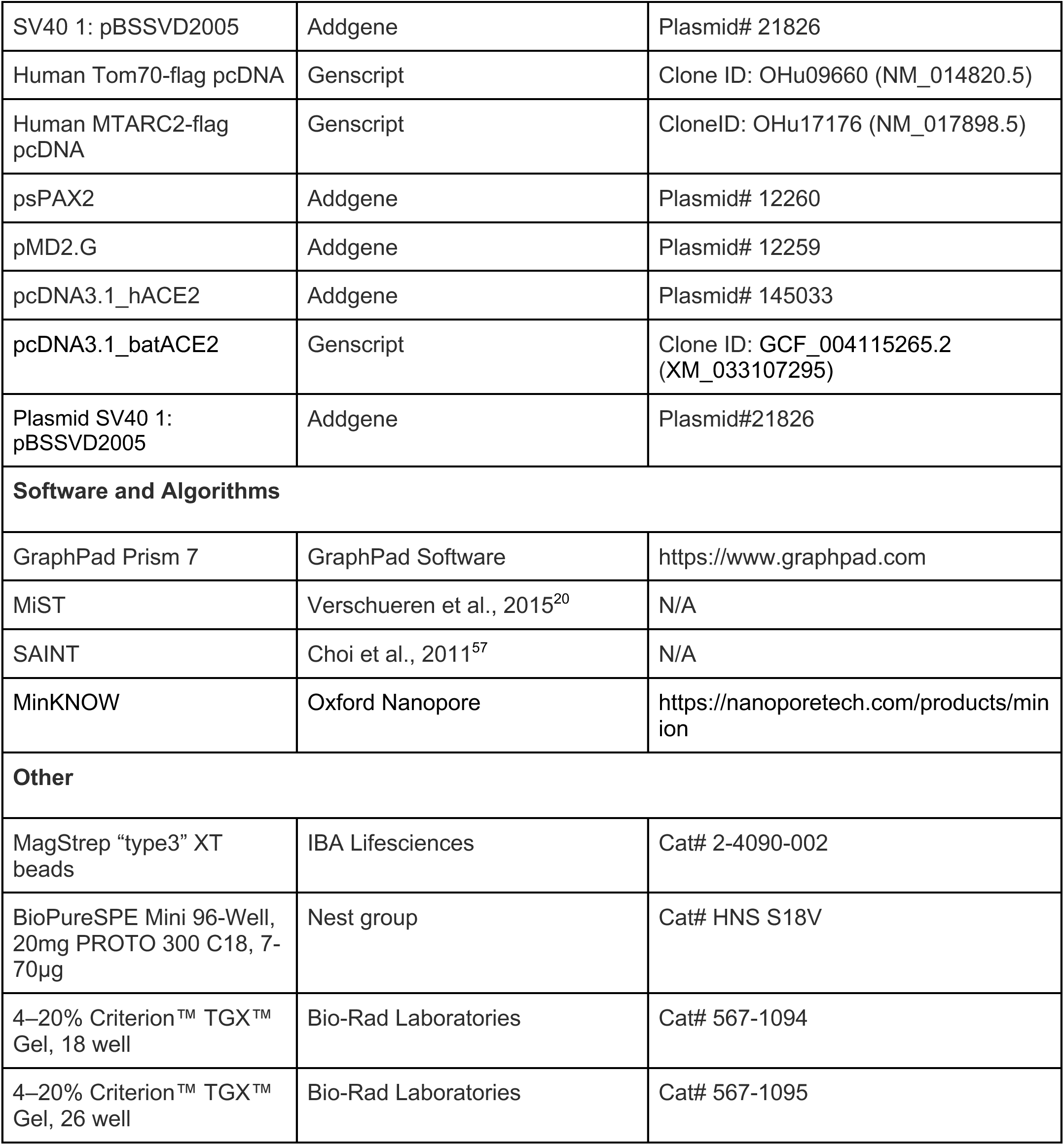

### Contact for Reagent and Resource Sharing

Further information and requests for resources and reagents should be directed to and will be fulfilled by the Lead Contact, Dr. Nevan J. Krogan.

## Experimental Details

### Capturing and sampling of *Rhinolophus ferrumequinum* bats

We sampled a colony of a few hundred individuals of the Greater horseshoe bat (*R. ferrumequinum*) occupying the tunnels of an abandoned water network system opening in a forest of evergreen oaks (Quercus ilex) with pastures near Jerez de la Frontera, Cádiz, Southern Spain in May 2020. In order to minimize disturbances to the colony, only an adult male bat and a pregnant female bat resting isolated from the group were captured with the help of a hand net. Following the guidelines of the American Society of Mammalogists (2016) and with the approval of the Spanish bio-ethical authority and the Andalusian regional government, the two bats were euthanized by cervical dislocation. Immediately after, bats were dissected and skin and lung samples together with the large bones (arms and legs) were preserved in a volume of 15 ml of RPMI medium in individual falcon tubes. Similarly, the fetus was beheaded and minced and preserved in RPMI medium. All samples were stored at 4 °C and processed in the lab within 25-30 hours of collection.

### Establishment of bat primary lung fibroblasts

Tissue samples from lungs were obtained as described above to establish bat primary cell cultures. Briefly, tissue fragments were incubated for 30 to 60 min at 37°C in a solution of 0.2 Wūnsch units/ml of Liberase (Sigma-Aldrich) in Dulbecco’s Modified Eagle Medium/Nutrient Mixture F-12 (DMEM/F-12) (Thermo Fisher) and 1% penicillin-streptomycin (10,000 U/mL) (Thermo Fisher). Digestion was stopped by adding DMEM/F- 12 supplemented with 15% fetal bovine serum (FBS) (Peak Serum) and 1% penicillin-streptomycin (10,000 U/mL). The tissue was mechanically dislodged by pipetting, followed by multiple centrifugations and washing steps. The pellet was resuspended and plated in a cell culture flask and incubated at 37°C in 5% CO2 for 5 to 7 days to allow cell migration from within the tissue fragments and attachment to the flask surface. Cells were then cultured in DMEM/F-12 medium supplemented with 10% FBS (Peak Serum) and 1% penicillin-streptomycin (10,000 U/mL) (Thermo Fisher).

### Immortalization of bat primary lung fibroblasts

Primary lung fibroblasts were immortalized by nucleofection with a SV40-LT antigen expressing construct (Addgene plasmid #21826, a gift from David Ron). Nucleofection was performed using P3 Primary Cell 4D- Nucleofector™ X Kit S following the protocol recommended for the transfection of primary cell lines. Briefly, 100,000 cells were resuspended in a nucleofection mix containing P3 Nucleofector solution (16.4µl), Supplement reagent (3.6µl) and SV40-LT plasmid (0.5µg). Electroshock was performed with Amaxa 4D- NucleofectorTM X Unit. Following nucleofection, cells were transferred into fresh culture medium.

### Development of *Rhinolophus ferrumequinum* lung fibroblasts stably expressing human ACE2 and TMPRSS2

*R. ferrumequinum* lung fibroblasts stably expressing human ACE2 and TMPRSS2 cells (RFe-AT) were obtained through lentiviral transduction. To generate lentiviruses, HEK293T cells were transfected with human ACE2 or TMPRSS2 lentiviral plasmids (Addgene #154981 and #170390, respectively), along with packaging plasmids for Gag-Pol (psPAX2) and VSV-G (pMD2.G) in a 15 cm dish. At 72 hours post-transfection, the supernatant was collected and filtered using a 0.45μm PVDF filter unit. Filtered virus supernatant was mixed with 8.5% PEG-6000 and 0.3M NaCl and incubated for at least 2 hours for precipitation. The virions were pelleted by centrifugation in a spinning bucket rotor at 3500 x g for 45 minutes at 4°C and resuspended in phosphate-buffered saline (PBS). For transduction, cells were trypsinized and spinoculated with hACE2 lentiviruses and 8μg/ml polybrene at 800 x g for 1 hour at room temperature. Cell pellet was then resuspended in fresh media and cultured for 96 hours. Cells were selected with 5μg/ml blasticidin for 3-4 days until non-transduced cells were dead. These cells were further spinoculated with hTMPRSS2 lentiviruses and transduced cells were selected with 250μg/ml hygromycin (Thermo Fisher).

### Cell culture

Human embryonic kidney cells with epithelial morphology (HEK293T/17) cells were maintained in Dulbecco’s Modified Eagle Medium (DMEM; Corning), supplemented with 10% fetal bovine serum (FBS; Gibco, Life Technologies) and 1% penicillin–streptomycin (Corning). Cells were cultured at 37 °C in a humidified incubator with 5% CO₂. The HEK293T/17 cell line was obtained from the UCSF Cell Culture Facility and cell line authentication was performed by the Berkeley Cell Culture Facility via STR profiling in August, 2017, confirming a 94% match with the reference HEK293T/17 profile. RFe cells were cultured in DMEM/F-12 medium supplemented with 10% FBS (Gibco, Life Technologies) and 1% penicillin-streptomycin (10,000 U/mL) (Thermo Fisher). Vero E6 (ATCC, CRL-1586) and TMPRSS2-Vero E6 (BPS Bioscience Cat# 78081) were maintained in Dulbecco’s modified Eagle’s medium (DMEM) (Corning) supplemented with 10% FBS (Peak Serum), 1% MEM non-essential amino acids solution (100x) (Gibco), 1% HEPES (N-2- hydroxyethylpiperazine-N-2-ethane sulfonic acid) (1M) (Gibco) and 1% penicillin/streptomycin (10,000 U/mL) (Corning) at 37 °C and 5% CO2. HEK293T (ATCC, CRL-3216), HEK293T AT (BEI Resources, NR-55293), and MRC5 (ATTC, CCL-171) were maintained in DMEM (Corning) supplemented with 10% fetal bovine serum (Peak Serum) and 1% penicillin/streptomycin (10,000 U/mL) (Corning) at 37 °C and 5% CO2. MRC5 hACE2 were generated for this study by transduction with lentiviruses applying the same protocol as for RFe- AT cells described above. All cell lines used in this study were regularly screened for Mycoplasma contamination, using the Universal Mycoplasma Detection Kit (ATCC, 30-1012K).

### Viruses and infections

Virus infections were performed using SARS-CoV-2, isolate USA-WA1/2020 (BEI Resources NR-52281). Additionally, four recombinant SARS-CoV-2 viruses, based on the USA-WA1/2020 reference sequence were used. The rSARS-CoV-2 WT have been previously described^58^. A recombinant virus with a single amino acid substitution in Nucleocapsid at position 80 (rSARS-CoV-2 N_P80T), a recombinant virus with two mutations leading to amino acid substitutions in Nucleocapsid at position 80 and in ORF9b at position 72 (rSARS-CoV- 2 N_P80T, ORF9b_T72I), recombinant virus with a single amino acid substitution in Nonstructural Protein 4 (NSP4_T269I), were generated for this study, using the same bacterial artificial chromosome (BAC)-based SARS-CoV-2 reverse genetic system previously described^44^. Briefly, two or one oligonucleotides were used to introduce N_P80T, ORF9b_T72I or NSP4T_269I coding changes into fragment 1 by site-directed mutagenesis. The region in the wild-type BAC between the unique restriction sites of BamHI and RsrII was replaced by the one from fragment 1 containing the mutation, and the newly generated BAC was used to produce the recombinant mutant viruses according to the protocol described previously. All viral stocks were grown in Vero E6 TMPRSS2 cells and validated by genome sequencing. Virus growth media (VGM) was used for all infections: Dulbecco’s modified Eagle’s medium (Corning) supplemented with 2% fetal bovine serum (Peak Serum), 1% non-essential amino acids (Gibco), 1% HEPES (Gibco) and 1% penicillin/streptomycin (Corning) at 37 °C and 5% CO2.

### Virus stock sequencing

Following viral RNA extraction using the Ribo Pure RNA Purification kit (Thermo Fisher Scientific) according to the manufacturer’s instructions, stocks were sequenced on the Oxford Nanopore MinION or Illumina MiSeq platforms. Samples sequenced on the Illumina MiSeq platform underwent cDNA synthesis using ProtoScript II (New England Biolabs, cat. E6560), followed by whole-genome amplification with two custom, interleaved tiling primer panels, as previously described^59^. PCR amplification was carried out using Q5 Hot Start High-Fidelity DNA polymerase (New England Biolabs, cat. M0493) in 25 μL reaction volumes. The resulting amplicons were purified using a 1.8X ratio of Ampure XT beads (Beckman Coulter, A63882) and used as input for library preparation using the Nextera XT kit (Illumina, cat. FC-131-1096) and sequenced in paired-end mode (2×150 nt) on the Illumina MiSeq. Genome assembly and quality control were performed using our custom vRAPID pipeline^60^. Complete genomes were classified by clade and lineage using Nextclade CLI (v3.15.3)^61^ and pangolin (v4.3)^62^ tools. For samples sequenced on an Oxford Nanopore flowcell, the Artic Consortium protocol^63^, was used with modifications. Native barcoding of amplicons was performed with Native Barcoding Kit 24 V14 (SQK-NBD114.24), according to the manufacturer’s instructions. Basecalling and demultiplexing was done with Dorado v0.9.6 (FAST5 input files) and v1.0.2 (POD5 input files) in high accuracy basecalling mode (dna_r10.4.1_e8.2_400bps_hac@v5.0.0). The genome assembly was done with the Artic pipeline (artic-network/artic-ncov2019, v.1.7.1)^64^ where reads were normalized to 10,000 then aligned to the reference genome Wuhan-Hu-1 (MN908947.3) using minimap2 (2.30-r1287), consensus variants were called with clair3 (v1.1.1). The consensus sequences were then analyzed for clade and lineage classification using the Nextclade CLI (v3.15.3)^60^ and pangolin (v4.3)^62^.

### SARS-CoV-2 adaptation to *R. ferrumequinum* lung fibroblasts

RFe-AT cells were seeded at the density of 2}10^5 cells/well on 12-well plate format. The next day cells were infected with SARS-CoV-2 WA1 isolate at an MOI of 1. At four days post-infection, 10µl of supernatant was passaged to a new culture of RFe-AT cells. A total of three blind passages were performed. Viral RNA was extracted from the supernatant of each passage using RiboPure kit (Invitrogen, AM1924) and subjected to whole virus genome nanopore sequencing.

### Generation of HIV-based pseudotyped particles

To generate HIV-based SARS-CoV-2 spike or VSV.G pseudotype particles, 5 × 10^6 293T cells were plated per 10-cm dish in 10 ml in growth medium. The next day, 7.5 µg pHIV-1NL4-3 ΔEnv luciferase reporter virus plasmid (NovoPro) and 2.5 µg SARS-CoV-2Δ19 plasmid (Invivogen) or pMD2.G (Takara) using LT-1 transfection reagent (Mirus) in DNA: reagent ratio 1:3 in 500µl of OptiMEM (Thermo Fisher) and added to cells drop-wise 20 min post incubation. At 12 hours post transfection cells were washed with PBS and 10ml of fresh medium was added. Supernatant was collected 48 hours post transfection, filtered through 0.45µm filters and aliquoted.

### Pseudotyped virus entry assay

To perform virus entry study 10^4 of RFe, RFe-AT, HEK293T and HEK293T-AT cells were seeded on 96- well plates. The next day cells were infected with HIV-based SARS-CoV-2 spike or VSV.G particles at MOI 1. At 48h post infection cells were lysed and firefly luciferase activity was assessed with ONE-Glo Luciferase Assay System kit (Promega). The read out was performed with a luminometer at a wavelength of 680nm.

### Infection of HEK293T cells overexpressing ACE2 orthologs with SARS-CoV-2 WA/01 virus

The day before transfection HEK293T cells were seeded in a 12-well plate format. The next day cells were transfected with 1µg of pcDNA3.1, pcDNA3.1 plasmid carrying human ACE2 or bat ACE2 with LT-1 reagent (Mirus). The pcDNA3.1 humanACE2 was a gift from Fang Li (Addgene plasmid # 145033; http://n2t.net/addgene:145033; RRID: Addgene_145033), a pcDNA3.1 plasmid carrying *R.ferrumequinum* ACE2 (batACE2) was designed and synthesized by Genewiz, based on the gene sequence available from the reference genome (NCBI RefSeq sequence: GCF_004115265.2; XM_033107295) with HA tag on its C-terminus. At 24h post transfection cells were infected with SARS-CoV-2 WA1/01 isolate at MOI 1. The next day cells were detached with PBS supplemented with 1mM EDTA (Gibco) and immunolabeled with hACE2 antibody (R&D systems, #AF933) conjugated with AlexaFluor647 (Thermo Fisher) and Fixable Viability Dye eFluor 450 (Thermo Fisher) for one hour on ice. Next cells were fixed in 4% paraformaldehyde for 15 minutes on ice, permeabilized with 1xPermWash buffer (BD Bioscience) and immunolabeled with anti-SARS-CoV-2 nucleocapsid (N) antibody conjugated to AlexaFluor488 (Thermo Fisher) for one hour on ice. Cells were washed with Cells Staining buffer (BioLegend) and resuspended in PBS prior cytometry analysis with Gallios cytometer (Beckman). At least 10,000 live cells were used to determine ACE2 expression level and infection rates for SARS-CoV-2 WA1 virus. Virus infection level in cells was measured by gating for SARS-CoV-2 nucleocapsid positive cell population.

### Immunofluorescence labeling of cells with α-tubulin antibody

Cells were seeded at a density of 3 }10^3 cells/well on 96 well plate format. The next day cells were fixed in 4% paraformaldehyde for 15 minutes, permeabilized with 0.1% Triton X-100 (Sigma-Aldrich) in PBS and blocked in 3% bovine serum albumin in PBS for 45 minutes. Next cells were incubated with mouse α-tubulin antibody (Cell Signaling) at a 1:200 dilution, followed by incubation with anti-mouse Alexa Fluor Plus 488 secondary antibody (Thermo Fisher Scientific) and DAPI (4’,6-diamidino-2-phenylindole) in 1:1000 dilution. All antibodies were diluted in 1% bovine serum albumin in PBS. Imaging was performed with EVOS M5000 microscope.

### Species origin characterization of RFe with RNAseq

RNA was extracted from four replicates using Qiagen RNeasy kits. Library preparation and sequencing were performed by Azenta Life Science using the following conditions: rRNA depletion for mRNA and long noncoding species, standard RNAseq run in Illumina HiSeq 4000 with a depth of 20-30 million reads per sample. The integrity of the RNA sequencing files of four samples (four replicates) was checked with the md5sum utility of GNU coreutils 8.22. For cleaning the reads and trimming, the software trim_galore v0.6.7^65^ was used, and to check the result of this process, FastQC v0.11.9^66^ and MultiQC v1.11^67^ were used. It was determined that the average quality throughout the sequence of cleaned reads was greater than a Phred Score of 35, so the analysis began. In order to obtain a robust result in verifying the species from which the samples came, it was decided to assemble the transcriptome of the four samples separately. For this purpose, the software Trinity v2.12.0^68^ was used. Once the transcriptomes of the four samples were assembled, the makeblastdb command of BLAST v2.13.0+ was used to build a database with each transcriptome. To verify the species of origin of the samples, the predicted mRNA sequences of the *R. ferrumequinum* cytochrome c and cytochrome b genes (XM_033088428.1 and XM_033112251.1 respectively) were used. The cytochrome c gene and the cytochrome b gene have been used in species identification studies previously^18,69^. With these sequences in FASTA format, searches were made with the blastn command of BLAST v2.13.0+ in the four databases generated with the transcriptomes, resulting in various contigs for each sample. To verify these results, the contigs identified by BLAST with the highest score and the lowest E-value for each gene and each sample were extracted from the assembled transcriptomes using seqtk^70^. With these contigs, the complete nt database of the NCBI BLAST website was searched to verify that the samples came from *R. ferrumequinum*.

### RaTG13 annotation and plasmid cloning

RaTG13 (MN996532.2) genome sequence was downloaded from GenBank and utilized to design 2xStrep- tagged expression constructs of ORFs and proteolytically mature Nsps derived from ORF1ab (with N-terminal methionines and stop codons added as necessary). SARS-CoV-2 and RaTG13 reading frames were codon-optimized for mammalian expression and cloned into pLVX-EF1alpha-IRES-Puro (Takara/Clontech) or pLVX-TetOne-Puro (Clontech) including a 5′ Kozak motif with either N-terminal or C-terminal 2XStrep tags using EcoRI and BamHI restriction sites. Similarly, human and bat host protein sequences were codon-optimized for *Homo sapiens* expression and inserted into pcDNA 3.1 vector (Addgene) with a C-terminal Flag-tag using HindIII and BamHI restriction sites. Restriction digestions were performed at 37°C for 1 hour, followed by T4 DNA ligase-mediated ligation at room temperature for 2 hours. The resulting constructs were transformed into chemically competent Stellar E. coli cells (Takara) for plasmid propagation.

### Transfection in HEK293T cells for AP-MS studies

HEK293T cells were transfected with RaTG13 viral protein constructs, along with a GFP control and a vector-only control (pLVX-EF1α-IRES-Puro), in three independent biological replicates for affinity purification. Ten million HEK293T cells were plated per 15-cm dish. Twenty-four hours later, cells were transfected with a total of 15 μg DNA per dish. The amount of bait plasmid used was based on prior SARS-CoV-2 affinity purification experiments^17^, considering relative expression levels and toxicity. Total DNA was adjusted to 15 μg with empty vector. The following amounts of bait plasmid were transfected per dish: 0.5 μg GFP; 3 μg Orf3a, Nsp14; 6 μg M; 7 μg Orf7b, Orf9c, Nsp5, Nsp6; 10 μg Nsp4, Nsp12, S; 15 μg N, Nsp1, Nsp2, Orf3b, Orf7a, Orf7b, Orf9b, Orf9c, Orf10, Nsp13, Nsp15, and empty vector. DNA was complexed with PolyJet Transfection Reagent (SignaGen Laboratories) at a 1:3 μg:μl ratio of DNA to transfection reagent, following the manufacturer’s instructions. After >40 h, cells were harvested using 10 ml DPBS without calcium or magnesium, supplemented with 10 mM EDTA for ∼5 minutes, followed by three washes with 10 ml DPBS. Each step was followed by centrifugation at 200 g for 5 min at 4 °C. Cell pellets were flash-frozen on dry ice and stored at −80 °C.

### Transduction in RFe cells for AP-MS studies

For lentiviral transduction, 0.5 million RFe cells were spinoculated with 400 μl of concentrated lentivirus in the presence of 8 μg/ml polybrene by centrifugation at 1,000 × g for 1 hour at room temperature. Transduced cells were then transferred to a T75 flask for recovery. Four days post-infection, cells were selected with 2.5 μg/ml puromycin until non-transduced control cells were completely dead. Following expansion, 10 million cells per 15-cm dish were seeded for affinity purification, with two dishes per sample in triplicate. The following day, doxycycline was added at 1 μg/ml to induce expression. Cells were harvested >24 hours after doxycycline induction. Cells were washed twice with cold DPBS, scraped in cold DPBS, and the suspensions were centrifuged at 200 × g for 5 minutes at 4 °C. Supernatants were removed, and cell pellets were snap-frozen on dry ice and stored at -80 °C.

### Sample preparation for AP-MS studies

Frozen cell pellets were thawed on ice for 10-15 minutes and resuspended in 1 ml IP lysis buffer (50 mM Tris-HCl, pH 7.4 at 4 °C; 150 mM NaCl; 1 mM EDTA), supplemented with 0.5% Nonidet P40 substitute (NP40; Fluka Analytical), cOmplete™ mini EDTA-free protease inhibitor cocktail, and PhosSTOP™ phosphatase inhibitor cocktail (Roche). Samples were incubated on a tube rotator at 4 °C for 30 minutes, frozen on dry ice for 20–25 minutes, partially thawed at 37 °C, and then fully thawed on ice. This freeze-thaw cycle was repeated once, followed by centrifugation at 13,000 g for 15 minutes at 4 °C to pellet cell debris. 40ul of Clarified cell lysates was mixed with SDS loading buffer for expression analysis by western blotting using the indicated antibodies. Up to 96 clarified lysates were arrayed into a 96-well deep-well plate for automated affinity purification using the KingFisher Flex (KFF) Purification System (Thermo Scientific) in the cold room. MagStrep ‘type3’ beads (40 μl; IBA Lifesciences) were equilibrated twice with 1 ml wash buffer (IP buffer with 0.05% NP40), and incubated with 0.95 ml lysate for 2 hours. Beads were washed three times with 1 ml wash buffer and once with 1 ml IP buffer. For on-bead digestion, beads were resuspended in 50 μl denaturation- reduction buffer (2 M urea, 50 mM Tris-HCl pH 8.0, 1 mM DTT), incubated at 37 °C for 45 minutes, and then brought to room temperature. Proteins were alkylated in the dark with 3 mM iodoacetamide for 45 minutes and quenched with 3 mM DTT for 10 minutes. Bead-bound proteins were digested overnight at 37 °C with 1 μl trypsin (0.5 μg/μl; Promega), followed by additional 0.5 μl trypsin for 2 hours. All incubation steps were performed with constant shaking at 1,100 rpm using a ThermoMixer C incubator. Resulting peptides were combined with 50 μl of 50 mM Tris-HCl (pH 8.0) to rinse the beads, then acidified with trifluoroacetic acid to a final concentration of 0.5% (pH < 2.0). Acidified peptides were desalted for mass spectrometry using a BioPureSPE Mini 96-Well Plate (20 mg PROTO 300 C18; The Nest Group) following standard protocols.

### MS data acquisition and peptide search

Samples were resuspended in 0.1% formic acid and analyzed using a Q Exactive Plus mass spectrometer (Thermo Fisher Scientific) coupled to an Easy-nLC 1200 ultra-high-pressure liquid chromatography (UHPLC) system via a Nanospray Flex nanoelectrospray ion source. Peptides were loaded onto a C18 reverse-phase column (25 cm × 75 μm, packed with ReproSil-Pur 1.9 μm C18 resin). Analytical columns were equilibrated with 5 μl of mobile phase A with a max pressure of 650 bar. HPLC buffer A consisted of 0.1% FA, and buffer B consisted of 0.1% FA / 80% ACN. Peptides were eluted by a linear gradient from 7 to 36% B over the course of 53 min at a flow rate of 300 nL/min, after which the column was washed with 95% B and re-equilibrated at 2% B. Each sample was analyzed with a 75-min acquisition, with all MS1 and MS2 spectra collected in the orbitrap in positive ion mode. MS1 scans were collected in profile mode with 70,000 resolving power, a 1}10^6^ AGC setting, 250ms maximum injection time, and a scan range of 300 - 1500 m/z. MS2 scans were collected in centroid mode with 17,500 resolving power, a 5}10^4^ AGC setting, 60ms maximum injection time, a 2.0 m/z isolation window, a normalized collision energy of 26, a loop count of 20 MS2 scans, a minimum AGC target of 8}10^3^, an intensity threshold of 1.3}10^5^, exclusion of unassigned and z=1 charge states, automated dynamic exclusion timing, and a scan range of 200 - 2000 m/z. Data was acquired using the Thermo software Xcalibur (4.2.47), Tune (2.11 QF1 Build 3006), and instrument suitability was monitored by QCloud^71^. For HEK293T samples, proteomic data were searched using MaxQuant (v1.6.12.0)^72^ against the UniProt-reviewed human proteome (downloaded February 28, 2020), enhanced green fluorescent protein (EGFP), and SARS-CoV-2 or RaTG13 protein sequences. Similarly, *Rhinolophus ferrumequinum* protein sequences were downloaded from UniProt and NCBI. To remove redundancy, sequences were grouped by Ensembl gene ID, and the longest sequence within each group was selected. Only sequences that appeared in both UniProt and NCBI were retained. RFe data were searched against this curated *Rhinolophus ferrumequinum* proteome, along with EGFP and SARS-CoV-2 or RaTG13 sequences. All raw MS data and search result files have been deposited in the ProteomeXchange Consortium via the PRIDE^56^ repository (PXD identifier PXD065873).

### Ortholog mapping of RFe proteins and percent identity

Orthologous gene pairs between *Rhinolophus ferrumequinum* and *Homo sapiens* were downloaded from Ensembl using the BioMart web interface (https://useast.ensembl.org/biomart/martview/b6f5f7374d580d6789a559a40e68a515). Ensembl gene identifiers were mapped to UniProt identifiers and corresponding protein sequences using UniProt ID mapping tables and the reference *H. sapiens* proteome. Orthologous protein sequences were aligned using the Needleman–Wunsch global alignment algorithm, implemented via the pairwiseAlignment function in the Biostrings R package (v2.52) with default parameters. Orthologous gene assignments were also compared against the InParanoid ortholog database^73^. Global alignments between *Rhinolophus ferrumequinum* and human proteins with the same gene name were performed using pairwiseAlignment from Biostrings (v2.68.1) in R. Sequence identity was calculated as 100 × (identical positions) / (length of shorter sequence). The percent identity between SARS-CoV-2 and RaTG13 viral proteins were calculated using Clustal Omega^74^.

### High-confidence protein interaction scoring and DIS analysis

Identified proteins were then subjected to PPI scoring with both SAINTexpress (version 3.6.3) and MiST (https://github.com/kroganlab/mist)^19,20^. High confidence PPIs were defined by a MiST score ≥ 0.7 and a SAINTexpress Bayesian false discovery rate (BFDR) ≤ 0.05. For all proteins meeting these thresholds, information on known stable protein complexes was retrieved from the CORUM database^75^. For a subset of viral baits, specifically RaTG13 M, Nsp12, Orf8, and Orf9b in HEK293T cells, and RaTG13 M in RFe cells, we observed a substantially higher number of PPIs passing the standard thresholds. For these baits, a more stringent filtering criteria was applied: AvgSpec > 3, MiST ≥ 0.8, and SAINTexpress BFDR ≤ 0.05. We then computed a Differential Interaction Score (DIS)^23^ for all interactions that passed the high-confidence criteria in at least one virus or species. The DIS was defined as the difference in interaction scores (K), calculated as the average of the MiST and SAINT scores, between the two viruses or species being compared. A DIS value near 0 indicates a shared interaction, while values approaching −1 or +1 indicate that the host–protein interaction is specific to one virus or species.

### Network generation and visualization

PPI) networks were generated in Cytoscape^76^ and annotated in Adobe Illustrator. Host–host physical interactions, protein complex definitions, and biological process groupings were obtained from CORUM^77^, Gene Ontology (biological process), and manually curated literature sources.

### Deep proteome profiling

The SP3 digest method was done similarly as described in Hughes et al ^78^. Cell pellets were thawed on ice and mixed with 1 ml of lysis buffer consisting of 50 mM HEPES pH 8 (Thermo Fisher Scientific #J63578AP), 1% (wt/vol) SDS (Sigma-Aldrich #L3771), 1% (vol/vol) Triton X-100 (Sigma-Aldrich #T9284), 1% (vol/vol) NP-40 (Sigma-Aldrich #74385), 1% (vol/vol) Tween 20 (Sigma-Aldrich #P7949), 1% (wt/vol) deoxycholate (Sigma-Aldrich #D6750), 5 mM EDTA (Corning #46-034-Cl), 50 mM NaCl (Sigma-Aldrich #S7653), 1% (vol/vol) glycerol (Sigma-Aldrich #15523), and 1 cOmplete protease inhibitor (Sigma #04693132001) (per 5 ml). Lysates were sonicated for 5s at 10% magnitude three times and centrifuged at max speed (13,000 rpm) for 5 minutes to pellet any insoluble material. The supernatant was then transferred to new Protein lo-bind tubes. Protein quantification was then performed by 660 assay (Thermo Fisher Scientific #22660) with ionic compatibility reagent (Thermo Fisher Scientific # 22663). 3.5 mg of each sample was transferred to new tubes. 100 mM Tris (2-carboxyethyl) phosphine (TCEP) (Thermo Fisher Scientific # 77720), 400 mM 2- chloroacetamide (Sigma-Aldrich #C0267) was added at 1:10 volume to each sample, and set in a Thermo Mixer to shake at 1,200 rpm at room temperature for 30 minutes. To prepare Sera-Mag bead solution for protein binding, SP3 beads (GE Healthcare, #45152105050250 and 65152105050250) were set at room temperature to warm. Beads were resuspended by pipetting, then equal parts of resuspended beads were combined into a slurry, with 350 µl bead solution added to each 3.5 mg reaction (4.5 ml of each bead type). Tubes were set on a magnetic rack and allowed beads to aggregate. Supernatant was removed and the beads were washed with 2x starting volume with water (700 µl per reaction, 18 ml total) three times. Supernatant was then removed, and the beads were resuspended at a concentration of 50 µg/µl. Beads were added to lysates at 1:10 weight of Sera-Mag bead solution to lysate (350 µl for 3.5 mg) to each, which were then resuspended by pipetting. Equal volumes of 100% ethanol were added to each sample for a final volume of 50% lysate, 50% ethanol. Tubes were set in a thermo mixer at 24°C for 10 minutes, shaking at 1,000 rpm. After mixing, samples were set in a magnetic rack, allowing beads to aggregate, after which the supernatant was removed and discarded. Protein bound beads were then washed three times with ∼1 ml 80% ethanol, removing all the supernatant between rounds. Beads were resuspended in 400 µl 50 mM Triethylamonium bicarbonate (TEAB) (Sigma-Aldrich # T7408). Trypsin (Promega # V5111) and lysC (Wako #125-02543) were added at a 1:100 ratio enzyme to protein. Samples were then set in a thermo mixer at 37°C shaking at 1,000 rpm overnight. The following morning, supernatant was transferred to a 15 ml falcon tube, and 300 µl washed Sera-Mag bead solution was added to each followed by 10 ml acetonitrile to induce peptide binding. Samples were vortexed and set in a magnetic rack. The supernatant was then discarded. Samples were washed with 5 ml 96% acetonitrile twice, then resuspended in 1 ml 96% acetonitrile and transferred to a 1.5 ml protein lo-bind tube. Samples were set in a magnetic rack, supernatant was removed and discarded. Samples were then fractionated as reported previously ^79^. Samples were sequentially eluted using isopropanol water gradients off the SP3 beads. Briefly, samples were resuspended in 800 µl 95% isopropanol, set in a magnetic rack and the supernatant transferred to a new protein lo-bind tube. This was then repeated with 90%, 85%, 80%, 75% and 70% isopropanol elutions into separate tubes. There was a final wash of water at the end to remove all remaining peptides from beads. Samples were then set in a speed-vac to dry down overnight. Prior to running on the mass spectrometer, samples were resuspended in 15 µl 2% acetonitrile, 0.1% formic acid and quantified by A205 with a 31 adjustment. Samples were adjusted to ∼500 ng/µl, with 1 µl run on the instrument column.

Digested samples were analyzed on an Orbitrap Exploris 480 mass spectrometry system (Thermo Fisher Scientific) equipped with an Easy nLC 1200 ultra-high pressure liquid chromatography system (Thermo Fisher Scientific) interfaced via a Nanospray Flex nanoelectrospray source. For all analyses, samples were injected on a C18 reverse phase column (25 cm × 75 μm packed with ReprosilPur 1.9-μm particles). Analytical columns were equilibrated with 6 μl of mobile phase A with a max pressure of 650 bar. Mobile phase A consisted of 0.1% formic acid (FA), and mobile phase B consisted of 0.1% FA / 80% acetonitrile (can). Peptides were separated by an organic gradient from 4% (2%) to 30% (25%) mobile phase B over 62 min followed by an increase to 45% (40%) B over 10 min, then held at 95% B for 8 min at a flow rate of 300 nl min−1. Data-independent analysis (DIA) was performed using an 80-minute gradient. An MS1 scan at 60,000 resolving power over a scan range of 350–1100 m/z, a normalized AGC target of 300%, and an RF lens setting of 40%. This was followed by DIA MS2 scans at 15,000 resolving power, using 20 m/z isolation windows over 350–1100 m/z at a normalized HCD collision energy of 30%. Loop control was set to All. Data was then searched using DirectDIA in Spectronaut v. 16.0.220606.53000 (Hawking) by searching against a database of Uniprot *Homo sapiens* sequences (downloaded 22 March 2022) and a *R. ferrumequinum* proteome using Biognosys (BGS) settings, variable modifications of methionine oxidation and N-terminal acetylation, and a fixed modification of carbamidomethyl on cysteine residues. Filtering was set to a final 1% false discovery rate (FDR) at the peptide, peptide spectrum match (PSM) and protein level, with no data normalization performed. Quantitative analysis was performed in the R statistical programming language (v.4.1.3). Initial quality control analyses, including inter-run clustering, correlations, principal component analysis (PCA), peptide and protein counts and intensities were completed with the R package artMS (v. 1.12.1). Statistical analysis of protein abundance changes between conditions were computed using peptide ion fragment data output from Spectronaut and processed using artMS. MSstats performs normalization by median equalization, no imputation of missing values, and median smoothing to combine intensities for multiple peptide ions or fragments into a single intensity for their protein group.

### Orf9b PPI and MSstats analysis

To quantitatively compare the PPIs of SARS-CoV-2 and RaTG13 Orf9b in HEK293T and RFe cells, we performed a separate AP-MS experiment. Cell pellets were collected for both Orf9b constructs and controls (empty vector and GFP) and processed for AP-MS as described above, with the exception that data was collected on an Exploris 480 mass spectrometer (Thermo Fisher Scientific) coupled to a Vanquish Neo ultra-high-pressure liquid chromatography (UHPLC) system via a Nanospray Flex nanoelectrospray ion source. Peptides were resuspended in 0.1% formic acid and loaded onto a PepSep C18 reverse-phase column (15 cm × 150 μm, packed with ReproSil-Pur 1.5 μm C18 resin). HPLC buffer A consisted of 0.1% FA, and buffer B consisted of 0.1% FA / 80% ACN. Peptides were eluted by a linear gradient from 2 to 28% B over the course of 32 min at a flow rate of 600 nL/min, after which buffer B was increased to 44% over the course of 18 min, and finally the column was washed with 90% B and re-equilibrated at 2% B. Each sample was analyzed with a 60-min acquisition, with all MS1 and MS2 spectra collected in the orbitrap in positive ion mode. MS1 scans were collected in profile mode with 120,000 resolving power, a 100% normalized AGC setting, automatic maximum injection time, and a scan range of 350 - 1250 m/z. MS2 scans were collected in profile mode with 15,000 resolving power, a 200% normalized AGC target, automated maximum injection time, a 1.3 m/z isolation window, a normalized collision energy of 30, exclusion of unassigned and inclusion of charge states z = 2-6, and dynamic exclusion within +/- 10ppm for 30 s after 2 observations. Data was acquired using the Tune (4.2 SP4). Proteomic data were searched using MaxQuant for both HEK293T and RFe samples. We used the artMS package (v1.18.0) in R to quantify changes in “prey” protein abundance between RaTG13 Orf9b WT and SARS2 Orf9b WT. The artmsQuantification function was used with default settings, except for MBimput = 0 and normalization_method = FALSE. All raw MS data and search result files have been deposited in the ProteomeXchange Consortium via the PRIDE repository^56^ (PXD identifier PXD065873).

### Network propagation

To investigate the connectivity between host proteins targeted by SARS-CoV-2 and RaTG13 proteins, we conducted a network propagation analysis. For the initial seed sets, we combined all identified prey proteins from our AP-MS experiments for each virus and propagated these seed nodes independently (16 total baits). The base network was the Reactome Functional Interactions (FI) network^80^. Network propagation was simulated using random walk with restart (RWR) to allow the initial signal from the seed proteins to spread across the network. A restart probability of 0.2 was used to balance exploration of the network while retaining influence from the initial seed nodes. We performed 20,000 simulations for each propagation to control for node degree bias. During each simulation, the labels of the seed proteins were randomly shuffled across the network to generate a null distribution. An empirical p-value was then calculated by determining the proportion of random propagation runs that produced a score greater than or equal to the observed propagation score for each node for the random propagations. Genes that were significant in either the SARS-CoV-2 or RaTG13 propagations were retained for downstream analysis. Additionally, to assess the connectivity of individual viral proteins, we performed separate network propagations for each SARS-CoV-2 and RaTG13 viral protein. For these analyses, the preys of each viral protein served as the initial seed set. Bait proteins with < 3 prey for either virus were excluded, resulting in the removal of 4 baits (Nsp1, Nsp5, Nsp14, S). The propagation was conducted the same as above with 20,000 simulations to account for network biases.

Gene set overrepresentation analysis (GSOA) was performed using the enricher function from the clusterProfiler package (version 3.12.0) in R with default parameters. For each analysis, significant GO terms (adjusted p-value < 0.05) were identified and refined to remove redundancy by constructing a GO term tree using pairwise distances calculated as 1 minus the Jaccard Similarity Coefficient of shared genes in GO Biological Process from MSigDB. The dendrogram was then cut at a height of 0.5 to define clusters of non-redundant gene sets. Within each cluster, the most significant term (i.e., lowest adjusted p-value) with the largest associated gene set was selected to ensure broad functional representation.

For the Orf9b subnetwork, we first isolated GO terms, and all associated genes, associated with “mitochondria” from the set of significant genes obtained from the initial, virus-wide, propagation results, found for either SARS-CoV-2 or RatG13. Using this gene list, we extracted a subnetwork of these genes from the Reactome FI network. This approach allowed us to focus on the functional relationships/interactions specific to mitochondrial pathways and evaluate their connectivity in the context of Orf9b across viruses.

### Co-immunoprecipitation assays

HEK293T cells were transfected with the mammalian expression constructs using Lipofectamine 2000 (Invitrogen) [ratio for lipo] according to the manufacturer’s instructions. Cells were harvested in Nonidet P-40 (NP-40) lysis buffer (50mM Tris-HCl pH 7.5, 150mM NaCl, 1mM EDTA, 0.5% NP-40) supplemented with cOmplete mini protease inhibitor tablets (Roche) and PhosSTOP phosphatase inhibitor tablets (Roche). Cell lysates were clarified at 13,000 RPM for 15 minutes at 4°C. Clarified cell lysates were subjected to immunoprecipitation with anti-strep magnetic beads (IBA) with end-over-end rotation for 2 hours at 4°C, followed by 5 washes with NP-40 buffer devoid of protease and phosphatase tablets. Protein complexes were eluted from the beads by direct incubation at 95°C in 1X SDS loading buffer. Eluates and clarified cell lysates were analyzed by immunoblotting with respective antibodies.

### Innate immune sensing assay in HEK293T cells

Each viral protein was overexpressed in HEK293T cells containing a Lucia reporter under the control of the IFN-β/ISG56 promoter, alongside an empty vector control. For viral protein expression, cells were transfected with 250 ng of either empty vector or a plasmid encoding Orf9b using Lipofectamine 2000 (Invitrogen). Twenty-four hours post-transfection, cells were stimulated with varying concentrations of poly I:C. After an additional 24 hours, 20 μL of culture supernatant from each well was transferred to a white 96-well opaque plate and mixed with 50 μL of QUANTI-Luc™ 4 Lucia/Gaussia “Glow” solution (InvivoGen). Lucia activity was measured using a SpectraMax iD3 plate reader, and fold induction was calculated relative to unstimulated control cells.

### RT-qPCR in RFe cells

RFe cells expressing Orf9b protein from SARS-CoV-2 or RaTG13, along with vector control were seeded in 12 well plates. Next day, cells were treated with 1 μg/ml doxycycline to induce expression. 24 h post-treatment, cells were infected with Sendai virus and were collected 24 h post-infection in the RLT buffer. RNA was extracted using the RNeasy Mini Kits (Qiagen) following manufacturer’s instructions Residual plasmid DNA was removed by on-column DNase digestion using RNase-free DNase (Qiagen), followed by cDNA synthesis using SuperScript™ IV First-Strand Synthesis System (Thermo Fisher). Real-time PCR was performed using the Fast SYBR Green Master Mix (Life technologies) on a CFX Opus Real-Time PCR system (Bio-Rad), with a PCR temperature profile as follows: 95°C for 20 seconds and then 40 cycles of 95°C for 3 seconds and 60°C for 30 seconds, followed by melt curve. The expression levels of RFe MX1, OAS1 and GAPDH mRNA were analyzed using gene specific primers as listed below. Expression levels were normalized to the GAPDH house-keeping gene and mRNA copies calculated relative to mock-infected cells. The following primers were used: RFe MX1: Fwd 5′-TTCCTCTGATTGTCCAGTTC-3′; Rev 5′- CCTCTTGTCGCTGGTGTCA-3′; RFe OAS1: Fwd 5′-GTTTGATCGCAGAGGAGAGTT-3′; Rev 5′- GTTCGCCCATCCATGATTCT-3′; RFe GAPDH: Fwd 5′-ATACCCACTCTTCCACCTTTG-3′; Rev 5′- CCTGTTGCTGTAGCCAAATTC-3′.

### RNAseq analysis

MRC5-ACE2 and RFe-AT cells were infected with rSARS-CoV-2 WT or mutants for 24 and 48h. RNA was extracted using the RNeasy Mini Kit (Qiagen) with on-column DNase digestion, following the manufacturer’s instructions. Library preparation and sequencing were performed by Azenta Life Science using the following conditions: polyA selection, standard RNAseq run in Illumina®, 2×150bp with a depth of 20 million reads per sample. Raw fastq files were downloaded from the Azenta server. Human (accession #GCF_000001405.40), Rhinolophus ferrumequinum (accession #GCF_004115265.2) and SARS-CoV-2 (accession #MN985325) reference genomes were downloaded from NCBI website. Read quality control, alignment and quantification were performed using the NF-core RNAseq pipeline [version 3.15]^81^. Default pipeline parameters were applied unless otherwise stated. Fastp [version 0.23.4]^82^ was used for adapter trimming, STAR [version 2.7.9a]^83^ for transcriptome alignment and Salmon [version 1.10.1]^84^ was used for quantification. To map reads to a given host and virus species simultaneously, host and viral reference fasta and gtf files were concatenated and used as input for processing. The generated transcript-level matrices were imported into R [version 4.2.3], and summarized to gene-level counts using tximport [version 1.26.1]^85^. Differential expression analysis was performed using the DEseq2 package [version 1.38.3]^86^. Briefly, the DEseq2 method models gene counts using a negative binomial distribution and applies the Wald test to identify genes that are differentially expressed among the groups. Genes with an absolute log2 Fold Change >= 1 & a BH-adjusted p-value < 0.05 were considered differentially expressed. To identify enriched pathways and functional gene-sets, we used custom R scripts utilizing clusterProfiler [version 4.6.2]^87^ and gene ontology (GO) gene sets and KEGG pathways for enrichment. R packages ggplot2 [version 3.5.2] and ComplexHeatmap [version 2.14.0]^88^ were used to visualize gene counts and differential analysis results. Selected ISGs were taken from a previous study^31^ and based on manual curation from the literature. All raw mRNA sequencing data files have been deposited in NCBI’s Gene Expression Omnibus^89^ and are accessible through GEO Series accession number: GSE302357.

### RaTG13 Orf9b-hTom70 structure modeling

To assess the structural impact of sequence variation on RaTG13 Orf9b, we performed comparative modeling of the RaTG13 Orf9b bound to Tom70 using MODELLER v10.1^90^. The experimentally determined structures of the human Tom70 bound to the N-terminal domain of the SARS-CoV-2 Orf9b (PDB IDs 7KDT and 7DHG) served as templates. We used FoldX^91^ to estimate the difference in binding affinity between the SARS-CoV-2 and RaTG13 structures.

### Orf9b evolutionary studies

To investigate Orf9b evolution, we assembled a diverse set of sarbecovirus genome sequences from multiple sources. This included 69 genomes from Sallard et al.^54^, 13 additional public genomes, and 59 unique genomes from the UCSC Genome Browser^92^. To identify additional informative sequences, we performed tblastn searches of the NCBI nr database using Orf9b from SARS-CoV-2 (Wuhan-Hu-1) and RaTG13, identifying three genomes with non-reference substitutions at positions 70 and/or 72: two bat viruses (Rp22DB159 and BtSY2) and one human SARS-CoV-2 variant (MW734954.1, labeled HuCoV2- LC0020256). To expand outgroup sampling, we conducted further tblastn searches using four divergent Orf9b sequences (from Bat_Hp-betaZhejiang2013, BtBM48-31, BtBtKY72, and BtRc-o319), retaining hits with ≥70% identity over ≥200 bp and lower similarity to RaTG13 or SARS-CoV-2 (Wuhan-Hu-1) than to the outgroup query. All collected genomes were aligned using MAFFT v7.526^93^, and the region corresponding to the annotated SARS-CoV-2 (Wuhan-Hu-1) N ORF was extracted from each genome. Candidate ORFs ≥800 bp were retained and used to construct an in-frame nucleotide alignment with MACSE v2.07^94^. Phylogenetic reconstruction was performed with PhyML v3.3.20200621^95^ using the GTR+F+I model and four rate categories. This analysis revealed a distinct clade in which an Orf9b-like overlapping ORF was conserved. We further filtered for unique N ORF sequences, reducing redundancy, and generated a final phylogeny rooted at the common ancestor of RaTG13 and BtBtKY72, following Sallard et al. Visualization of the tree and corresponding alignments was performed using the ggtree^96^ and ggmsa^97^ R packages. We noted that coronavirus evolutionary history is complicated by numerous recombination events^54^. While we made the tree from the N ORF, and show alignments for regions within N ORF, recombination means that this single phylogeny does not fully reflect the history of these genomes.

### SARS-CoV-2 mutation frequency

We obtained time-resolved mutation frequencies of the Orf9b T72I mutation (C28498T) in human SARS-CoV-2 isolates from the CoV-Spectrum site^55^, which uses data from GISAID^98^.

## Acknowledgements

We would like to thank John Huddleston and Katie Kistler for useful discussions on SARS-CoV2 mutation frequencies. We also thank Richard Cadagan for excellent technical assistance and Dr. Randy Albrecht for support with the BSL3 facility and procedures at the <u>ISMMS.</u> We thank Dr. Arturo Marin for his previous bioinformatics analysis for this work, conducted as part of his postdoctoral fellowship from December 1, 2020 to December 14, 2023, at the Icahn School of Medicine at Mount Sinai. We thank Delini Samarasignhe for her assistance in collecting cell pellets for AP-MS studies in bat cells. This research was funded by grants from the National Institutes of Health (U19AI135990 to N.J.K, M.O., J.B.; U19AI135972 to A.G-S., N.J.K, J.B.; U54AI170792 to H.S.M). H.S.M. is an Investigator of the Howard Hughes Medical Institute. I.P.C. was supported by NIH/NIAID (F31 AI164671-01). M.O. received support from NIH U19 AI135990, the James B. Pendleton Charitable Trust, Roddenberry Foundation, P. and E. Taft, and the Gladstone Institutes. M.O. thanks Fast Grants and the Innovative Genomics Institute for their support. M.O. is a Chan Zuckerberg Biohub – San Francisco Investigator. Experiments performed in the NJ laboratory were funded by ANR EmerCoV AAP CE35. Research reported in this publication in the BSL3 facility was supported by the NIAID/NIH under Award Number G20AI174733 (R.A. Albrecht). This work was supported in part through the computational and data resources and staff expertise provided by Scientific Computing and Data at the Icahn School of Medicine at Mount Sinai and supported by the Clinical and Translational Science Awards (CTSA) grant UL1TR004419 from the National Center for Advancing Translational Sciences. Research reported in this publication was also supported by the Office of Research Infrastructure of the National Institutes of Health under award number S10OD026880 and S10OD030463. The content is solely the responsibility of the authors and does not necessarily represent the official views of the National Institutes of Health.

## Author contributions

The following authors designed and conceptualized the study: NJK, AGS, LM, JB, HSM. The following authors performed experiments or data acquisition: JB, MR, CY, RA, DA, SV, AM, SM, AC, TK, JMM, AR, SM, MM, HF, IPC, CD, NJ, SG, ICV, AB, MDS, SM, MM, HF, CSF, SG, RS TYT, IM, SA, JJ, CI, CMR. The following authors conducted data processing or analysis: JB, MR, YZ, MG, BP, IPC, JMY, DMW, IE, DTC, RF, CD, TYT, ME, ZK, IM. The following authors supervised research: NJK, AGS, LM, LMS, MO, DLS, HVB, BP, MB, TH, HSM, JSF, KAV, RMS, NJ, KO, LZA. The following authors drafted the original manuscript: JB, MR, ME. All authors edited the manuscript.

## Declaration of generative AI and AI-assisted technologies in the writing process

During the preparation of this manuscript, the authors used ChatGPT to rephrase or shorten portions of the text. All content generated with the assistance of this tool was subsequently reviewed and edited as needed by the authors, who take full responsibility for the content of the publication.

## Supplementary Figures legends

**Figure S1. Characterization of immortalized *R. ferrumequinum* lung fibroblast cell lines (RFe).**

(A) Heatmap showing relative expression of fibroblast-associated markers in bat (RFe) and human (MRC5) fibroblasts compared to HEK293T cells, based on global proteomic profiling. Values represent log₂ fold-change (log₂FC) relative to HEK293T; black boxes indicate proteins detected in only one condition.

(B) Representative immunofluorescence images showing cellular morphology of RFe and MRC5 fibroblasts. Cells were stained with anti-α-tubulin (green) and DAPI (blue) to visualize the cytoskeleton and nuclei, respectively. Scale bars, 100 µm.

**Figure S2. Comparative analysis of viral protein expression, and host interaction profiles of SARS-CoV-2 and RaTG13.**

(A) Percent amino acid identity between SARS-CoV-2 and RaTG13 proteins. Proteins that are 100% identical are indicated in black.

(B) Expression levels of individual viral proteins used as baits in AP-MS experiments. Bar plots show log₂ intensity values measured by mass spectrometry in HEK293T (top) and RFe (bottom) cells. Color indicates the virus type: SARS-CoV-2 (green), RaTG13 (purple), and shared (gray).

(C-E) Immunoblot detection of 2XStrep-tagged RaTG13 in HEK293T cells (C) and RaTG13 (D) or SARS-CoV-2 (E) proteins in RFe cells. One representative input lysate per bait (from n=3 replicates used for AP-MS) was probed with anti-Strep antibody. Arrowheads indicate the expected bands corresponding to each viral protein.

(F) Venn diagram showing overlap in high-confidence PPIs of SARS-CoV-2 and RaTG13 in HEK293T or RFe cells (top); and PPIs of SARS-CoV-2 and RaTG13 across the two cell types (HEK293T vs. RFe). Comparisons across viruses exclude PPIs involving identical baits between SARS-CoV-2 and RaTG13.

**Figure S3. Conserved and species-specific host interaction landscapes of RaTG13 and SARS-CoV- 2.**

(A) Comparative differential interaction map of RaTG13 in bat (RFe) versus human (HEK293T) cells. Edge color represents the differential interaction score (DIS): blue edges indicate interactions enriched in human cells, red edges indicate those enriched in bat cells, and grey edges denote shared interactions. Each bait node is split into two halves, representing the log₂ intensity values of bait expression in human (left) and bat (right) cells. Thin grey edges indicate known host-host (human-human) physical interactions. Host proteins belonging to the same protein complex or biological process are shaded in light yellow and light blue, respectively. Host-host interactions, complex annotations, and biological process groupings were derived from CORUM, Gene Ontology (biological process), and manual curation.

(B) Comparative differential interaction map of SARS-CoV-2 in bat (RFe) versus human (HEK293T) cells.

The PPIs for identical baits between SARS-CoV-2 and RaTG13 are shown only in the SARS-CoV-2 PPI map.

**Figure S4. Shared and bat-specific host interactions of SARS-CoV-2 and RaTG13 viral proteins.**

(A) Heatmap depicting bat-specific PPIs for SARS-CoV-2 and RaTG13. The left panel shows the log₂ fold change in host protein abundance in RFe compared to HEK293T cells (RFe/HEK), and the right panel shows the K score (the average of MiST and SAINT scores) of viral-host PPIs identified specifically in bat cells. Asterisks denote proteins not detected by mass spectrometry of whole cell lysates.

(B) Heatmap showing host proteins that interact with SARS-CoV-2 and RaTG13 viral proteins in both bat (RFe) and human (HEK293T) cells. Only interactions with an interaction score (K score) > 0.5 in all four datasets are shown. Host proteins are annotated by functional categories or known complexes.

**Figure S5. Network propagation analysis identifies mitochondrial pathway signature of SARS-CoV-2.**

(A) Comparative significance scatter plot showing -log₁₀(p-values) for genes identified via network propagation seeded with host protein interactors of SARS-CoV-2 (y-axis) and RaTG13 (x-axis). Genes significantly enriched only in the RaTG13 or SARS-CoV-2 networks are shown in purple and green, respectively, while those enriched in both are shown in black. Numbers in each quadrant indicate the number of significant genes after network propagation analysis relative to 20,000 random shuffling of node labels, including those identified from PPI data and through network propagation.

(B) Gene Ontology (GO) enrichment analysis of significant propagated genes (adj. p-value ≤0.05) from each quadrant in A. Numbers within cells indicate the number of genes per category, with color intensity corresponding to –log₁₀(adj. p-value).

(C) Scatter plot showing the number of mitochondrial-related GO terms of significant propagated genes of each of the viral proteins from RaTG13 (x-axis) and SARS-CoV-2 (y-axis).

(D) Heatmap showing the number of genes associated with mitochondria-related GO terms from network propagation analysis for each viral protein.

(E) Subnetwork of SARS-CoV-2 Orf9b interactions with mitochondrial proteins. Network edges reflect interactions from AP-MS (black) or the Reactome database (green: TOM complex; blue: TIM complex), filtered for nodes annotated in mitochondrial GO pathways.

**Figure S6. Evolutionary and secondary structural context of Orf9b.**

(A) Secondary structure prediction of SARS-CoV-2 and RaTG13 Orf9b using JPred 4 server. Red cylinders indicate predicted α-helices, and the amino acid sequence from residues 60-80 is shown above.

(B) Phylogenetic tree of 74 diverse sarbecovirus genomes, constructed using nucleotide alignments of the N open reading frame and rooted at the common ancestor of RaTG13 and BtKY72 (as in Sallard *et al*.^54^). The aligned region encoding Orf9b positions Q70 and T72 is shown, with N-frame amino acid translation (left), nucleotide sequence (middle), and Orf9b-frame translation (right).

(C) Time-resolved mutation frequency of the Orf9b T72I mutation (nucleotide change C28498T) in circulating SARS-CoV-2 isolates, obtained from CoV-Spectrum^55^.

**Figure S7. Transcriptomic profiling reveals enhanced ISG induction by Orf9b T72I mutant virus.**

(A) Gene Ontology (GO) enrichment analysis of genes upregulated in MRC5 cells infected with the Orf9b_T72I, N_P80T mutant virus compared to N_P80T virus at 24 hpi. Color intensity represents -log₁₀(adj. p-value), and the number in each box indicates the number of genes per term.

(B) Representative expression plots of selected ISGs from RNA-seq in human (top row) and bat (bottom row) cells.Log₂ fold-change values relative to mock-infected controls are shown at 24 and 48 hours post-infection (hpi) for each gene. Cells were infected with either N_P80T or Orf9b_T72I/N_P80T virus, with the T72I mutant consistently showing higher ISG induction over time.

## Table legends

**Table S1. Summary of BLAST results for the hypothetical cytochrome b sequence queried against the complete nucleotide database.** Shown are results from uninfected samples at time 0 (UI-0h), replicates 1-4. In all cases, the top BLAST hit corresponds to the cytochrome b sequence of *Rhinolophus ferrumequinum*.

**Table S2. Summary of BLAST results for the hypothetical cytochrome c sequence queried against the complete nucleotide database.** The BLAST results for uninfected samples at time 0 (UI-0h) for replicas 1 to 4 are shown. The top BLAST hit for sample UI-0h-2 was *Rhinolophus ferrumequinum*. For samples UI-0h-1, UI-0h-3, and UI-0h-4, the top hit was *Hipposideros armiger*, with *Rhinolophus ferrumequinum* as the second-best match in all cases (see Supplementary Material).

**Table S3: List of high-confidence PPIs of SARS-CoV-2 and RaTG13 in RFe and HEK293T cells.** Spectral counts, (column AvgSpec), SAINT BFDR and MiST scores are shown for each dataset. RaTG13 viral protein sequences are also listed.

**Table S4. Percent protein identity between *Rhinolophus* and human orthologs.** Percent identity is shown for each high-confidence prey identified across all four datasets, comparing human and *Rhinolophus ferrumequinum* orthologous proteins.

**Table S5. Comparative analysis of Orf9b PPIs between RaTG13 and SARS-CoV-2 in bat and human cells.** The table lists the log₂ fold change and adjusted p-values (adj p) for each PPI, calculated using MSstats. Positive values indicate stronger enrichment with RaTG13 Orf9b, while negative values indicate stronger enrichment with SARS-CoV-2 Orf9b. Data are shown separately for RFe and HEK293T cells.

**Table S6. Differential expression of interferon-stimulated genes (ISGs) in RFe-AT and MRC5-ACE2 cells infected with SARS-CoV-2 N_P80T or Orf9b_T72I, N_P80T mutant viruses.** Log₂ fold change and adjusted p-values from RNA-seq analysis are shown for ISGs at 24 and 48 hours post-infection in both cell types.

