## Supplementary figures and images for "Coronavirus protein interaction mapping in bat and human cells identifies molecular and genetic switches for immune evasion and replication"

### Figure S1

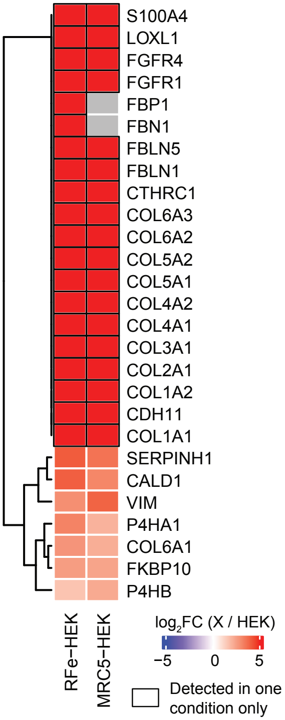

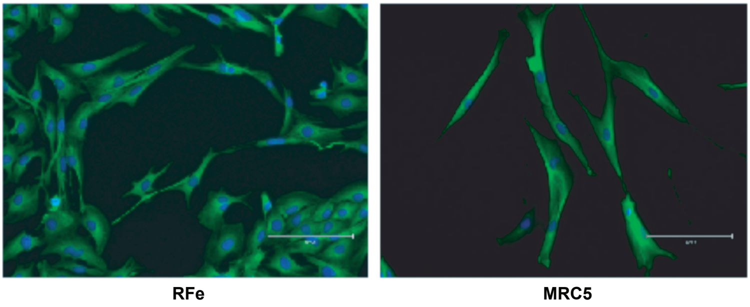

### Figure S2

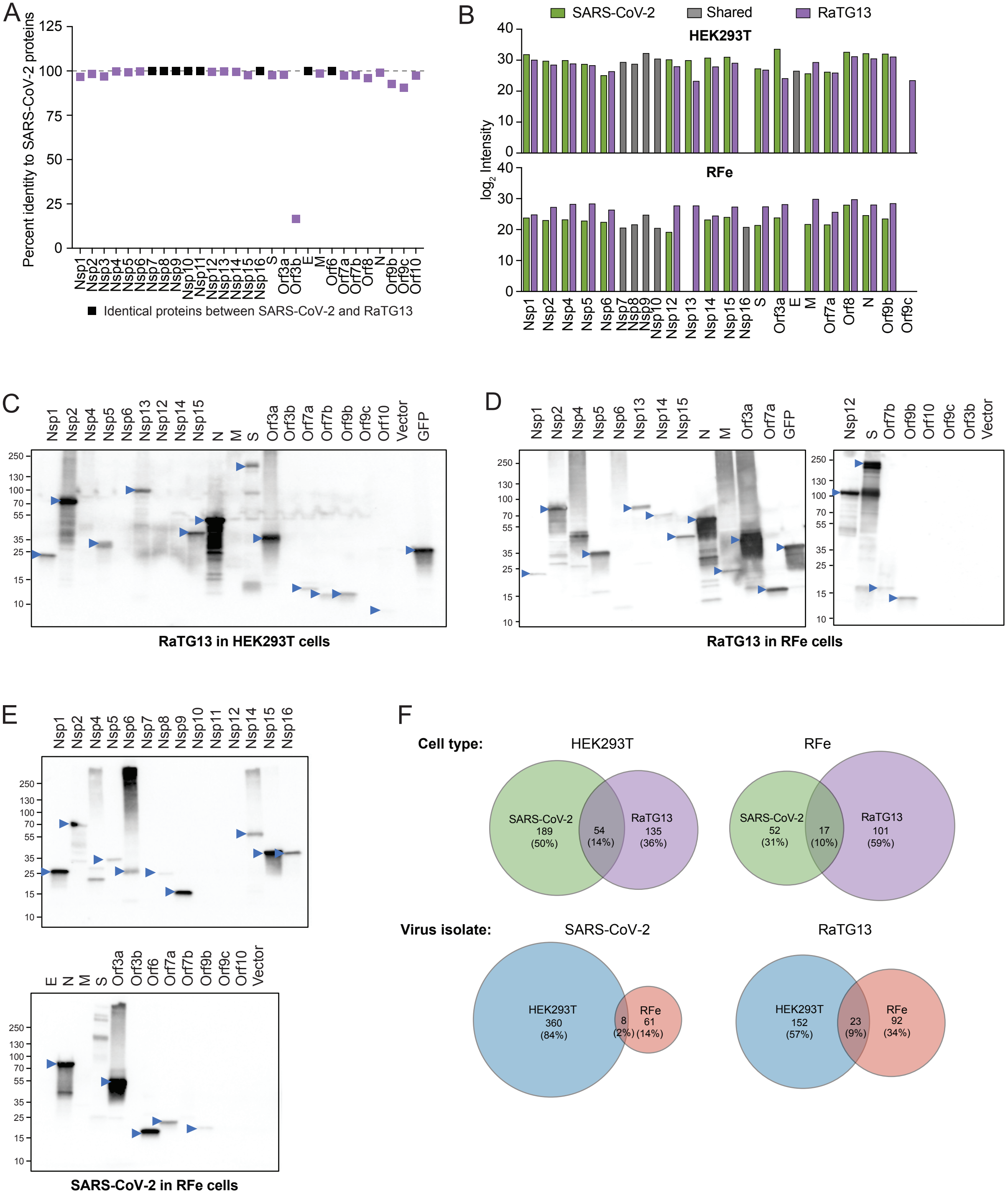

### Figure S3

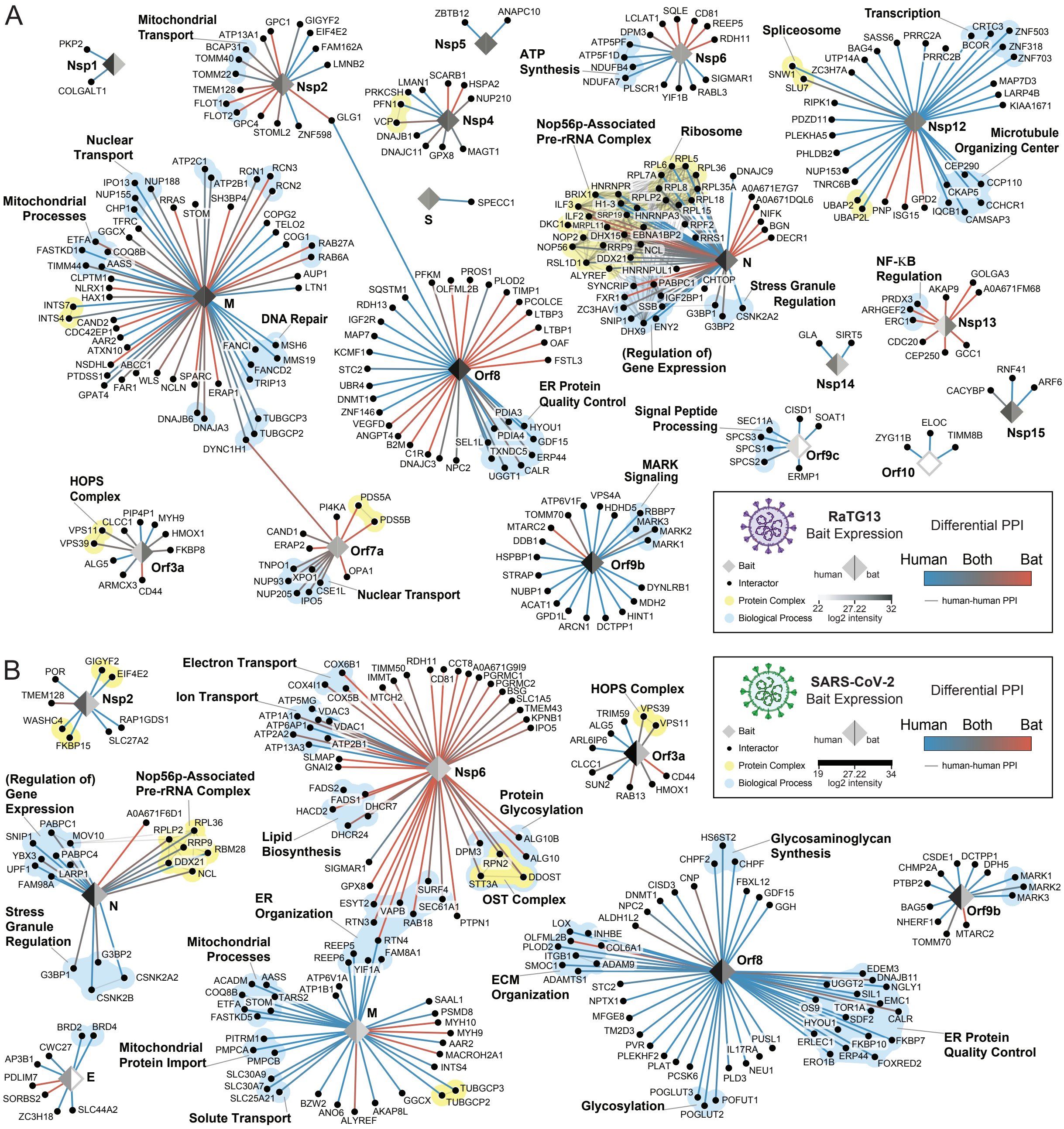

### Figure S4

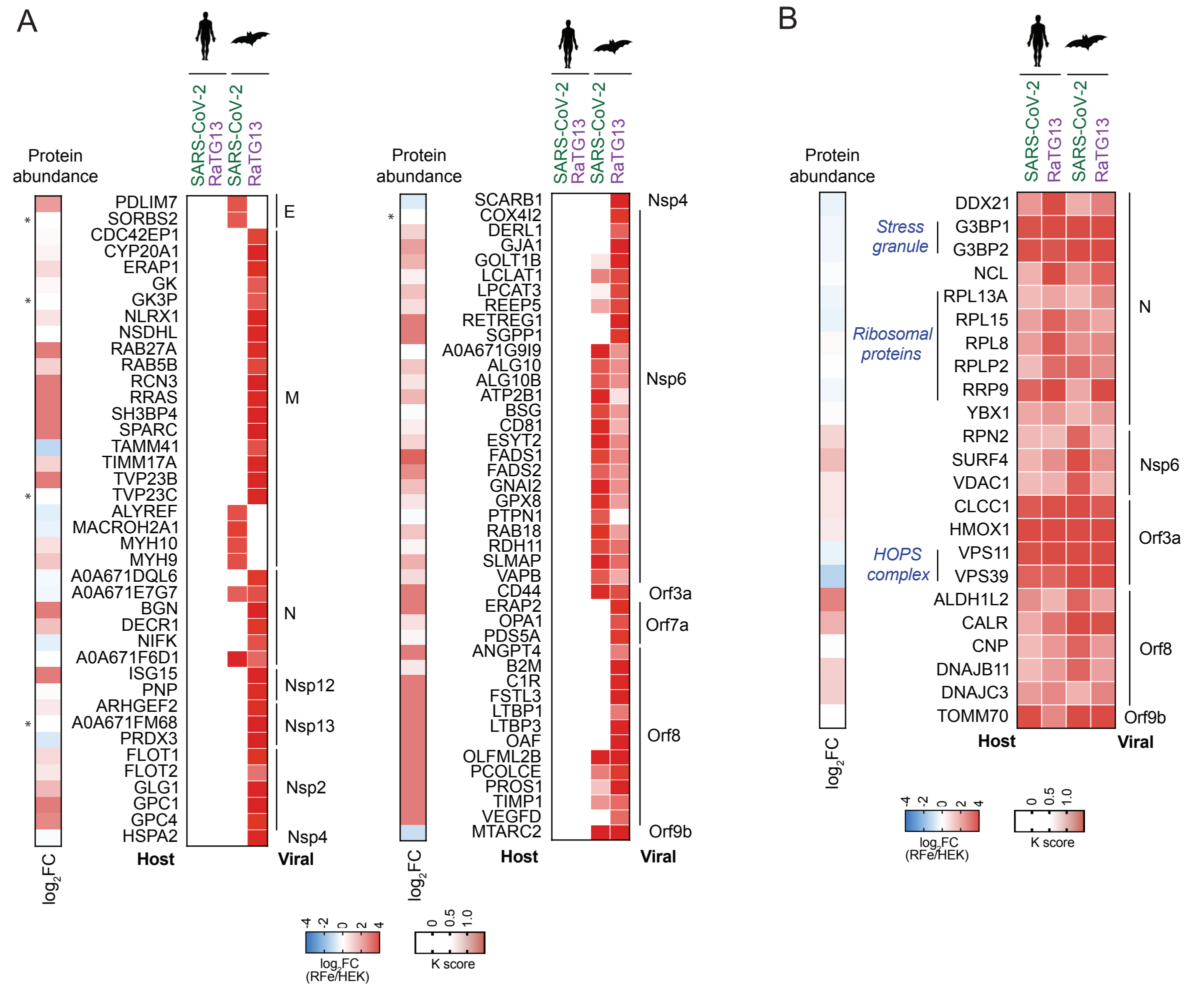

### Figure S5

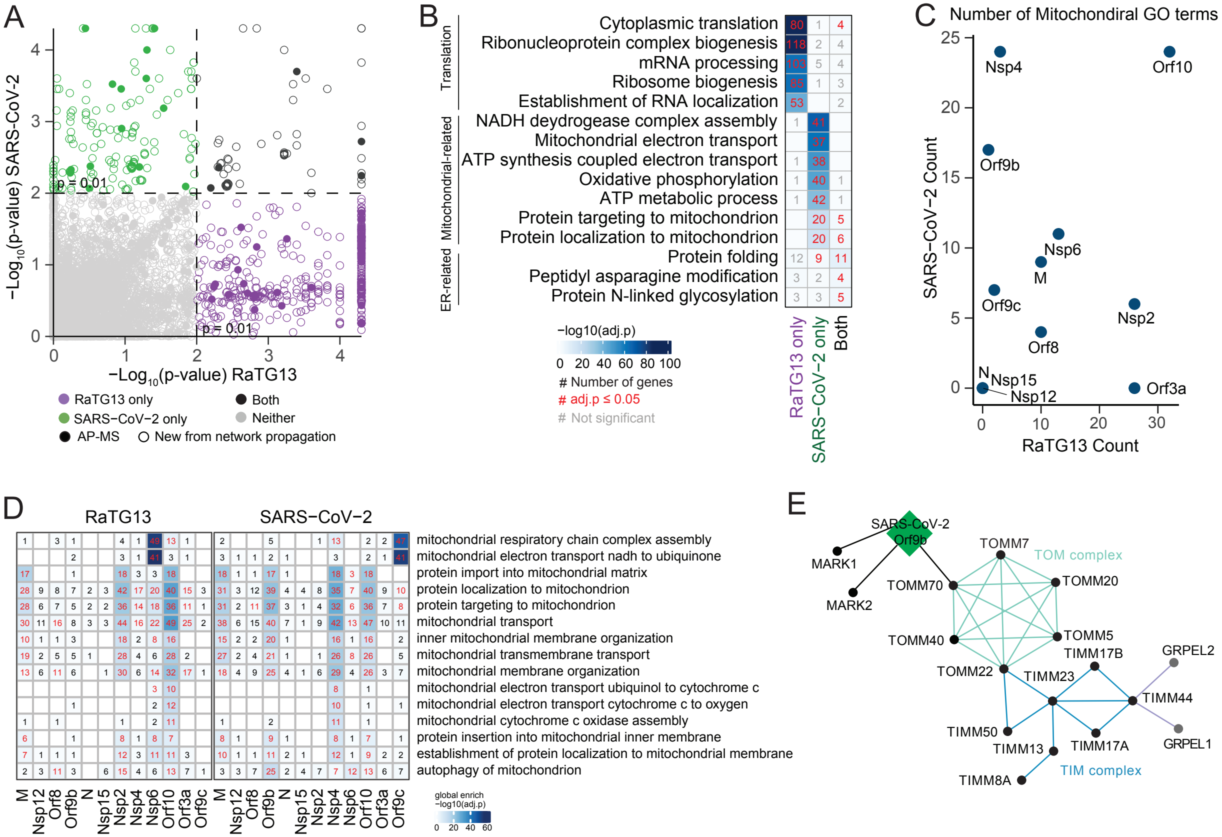

### Figure S6

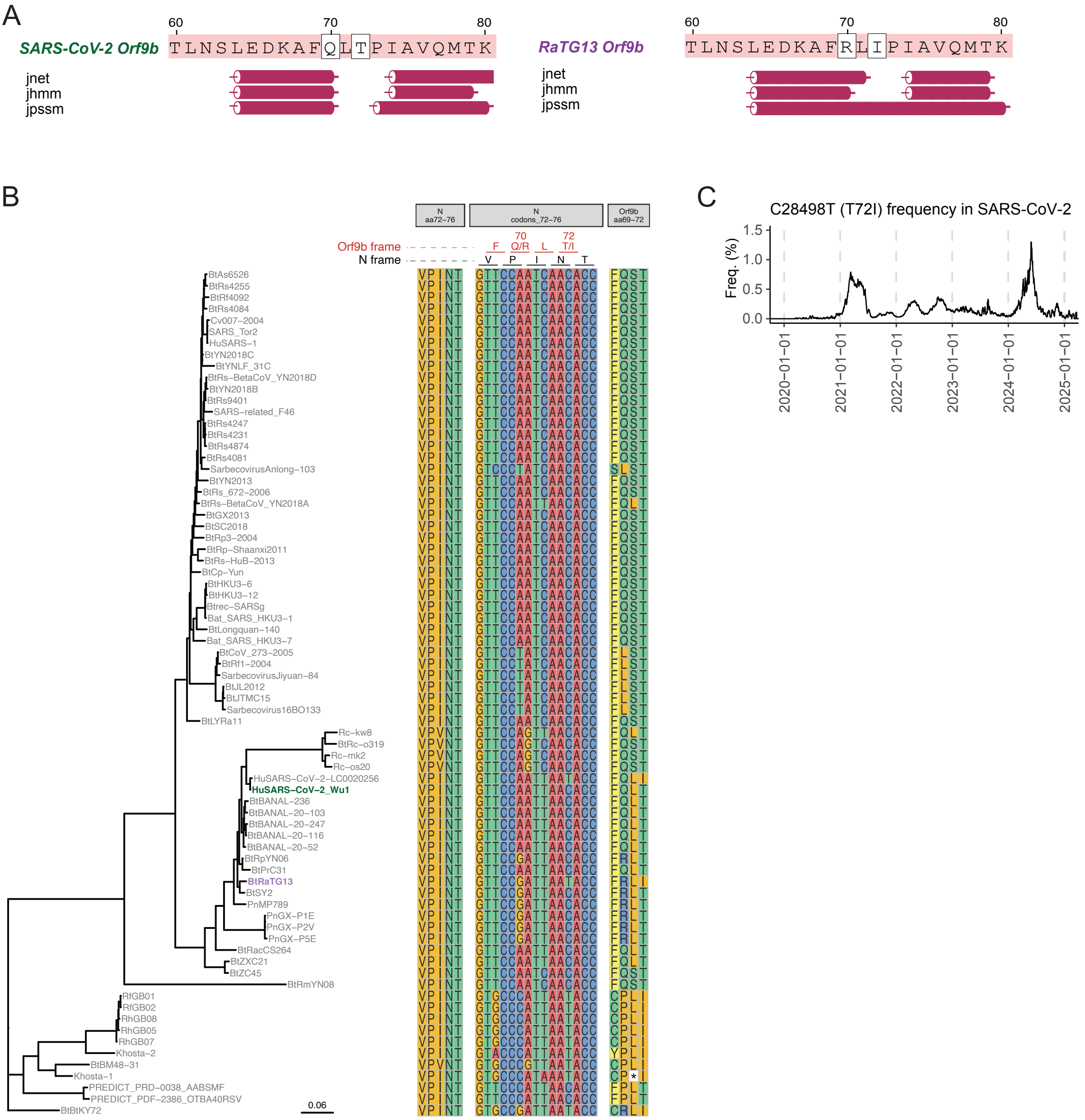

### Figure S7

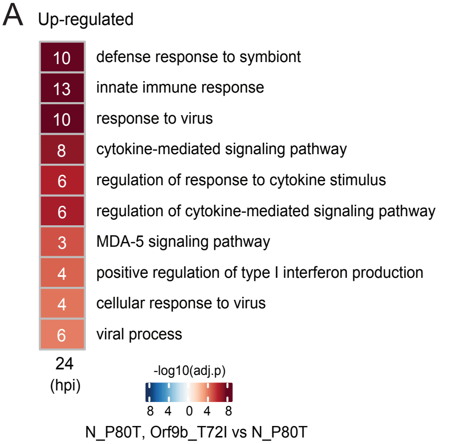

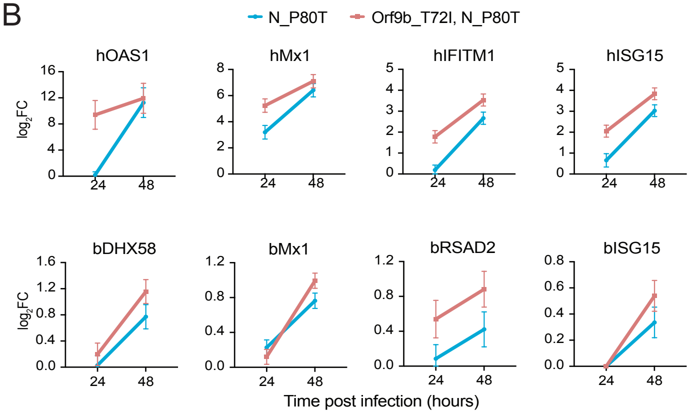
